# Sirtuin 6 is a histone delactylase

**DOI:** 10.1101/2024.09.28.615627

**Authors:** Garrison A. Nickel, Nicholas J. Pederson, Faheem, Zhenyu Yang, Jack Bulf, Katharine L. Diehl

## Abstract

Histone lactylation (Kla) is a post translational modification (PTM) that is derived from metabolic lactate. Histone Kla has been extensively studied in the field of inflammation resolution and macrophage polarization but has also been implicated in diverse cellular processes including differentiation, various wound repair phenotypes, and oncogenesis in several cancer models. While mechanistic connections between histone Kla and transcriptional changes have been studied in very limited contexts, general mechanistic details describing how regulation of gene expression by histone Kla occurs are scarce. It is hypothesized that histone Kla may be installed either through nonenzymatic means or by the p300 histone acyltransferase complex, and it is known that Class I HDACs and Sirtuins 1-3 can remove histone Kla. Here, we identified histone delactylase activity of the deacylase enzyme Sirtuin 6 (Sirt6), a member of the Class III HDAC family known to have roles in regulating metabolic homeostasis. We characterized the ability of Sirt6 to delactylate histones *in vitro* and in mammalian cell culture. We identified H3K9 and H3K18, canonical histone sites of Sirt6-catalyzed deacetylase activity, as sites of its delactylase activity. We also demonstrated that the delactylase activity of Sirt6 and the Class I HDACs are additive, suggesting that they represent different cellular axes of regulating gene expression via controlling levels of histone Kla.

## Introduction

Histones, the primary protein component of eukaryotic chromatin organization, are heavily modified with a wide variety of post-translational modifications (PTMs).^1,2^ While PTMs such as histone lysine methylation (Kme) and lysine acetylation (Kac) have been extensively studied and their roles are relatively well-defined,^3–6^ in recent years mass spectrometric approaches have identified a host of novel histone PTMs which have not yet been fully characterized. These newly discovered PTMs, such as lysine β-hydroxybutyrylation^7,8^, succinylation^9–11^, malonylation,^10^ and propionylation,^12,13^ incorporate diverse primary metabolites into histone modifications. One prevalent hypothesis posits that these metabolite-derived PTMs are indicative of a regulatory axis through which high levels of metabolites can be ‘sensed’ by the cell and converted into a transcriptional response.^14–16^

Originally discovered in 2019,^17^ histone lysine lactylation (Kla) is a leading example of how cellular primary metabolite pools can influence gene expression by being directly incorporated into histone PTMs. Measurable increases in histone Kla result from treatment of cells with supplemental lactate or under conditions which bias metabolism towards glycolysis and lactate production.^17,18^ Isotopic labelling experiments have demonstrated that extracellular lactate and extracellular glucose are both incorporated into Kla modifications, directly linking cellular metabolism to histone modifications.^17^ There is some debate about the immediate source of histone lactylation, with some studies pointing to glyoxalate metabolism as a source for reactive lactyl moieties.^19,20^ However, a recent study found that L-lactylation is the dominant stereoisomer of histone Kla present in cells, implicating lactyl-CoA as the precursor to histone lactylation.^21^ The concentration of lactyl-CoA detected in HepG2 cells under typical culture conditions (i.e., no lactate supplementation, high glucose) is about 1,000 times lower than of acetyl-CoA, but recent studies have found that lactyl-CoA levels are highly sensitive to glucose metabolism in these same cells.^21,22^ A body of recent experimental evidence implicates histone Kla in a variety of biological processes and diseases, including oncogenesis and cancer progression,^23–28^ differentiation,^29^ anti-inflammatory response,^17,18,30–32^ tissue repair,^33^ angiogenesis,^34,35^ and metabolic diseases such as diabetes mellitus.^36,37^ However, major outstanding questions remain: to what extent is Kla functionally distinct from Kac and how does this distinction occur. Many studies that report changes in levels of histone Kla also report similar changes in levels of Kac,^17,38^ making it difficult to disentangle the roles of the two PTMs. Additionally, since conditions typically used to stimulate histone Kla do so by increasing lactate levels, it is difficult to deconvolute the effects of histone Kla from other signaling effects caused by extracellular and/or intracellular lactate.^38,39^

By identifying the enzymes that regulate installation or removal of Kla, it may be possible to selectively increase or decrease cellular Kla levels without affecting cellular lactate concentrations, histone Kac, and/or other histone PTMs, thus allowing the downstream effects of histone Kla to be independently measured. While it is still not completely clear how histone lactylation is installed, some experimental evidence indicates that it may be deposited by the p300 histone acyltransferase complex, the enzyme HBO1, or by a nonenzymatic pathway involving lactylglutathione.^17,19,20,40^ Several mechanisms by which histone Kla removal occurs have been described. Moreno-Yruela et al. showed that the class I histone deacetylases (HDACs) are able to remove histone Kla, although they also deacylated all other PTM substrates tested, exhibiting no obvious substrate selectivity.^41^ The Sirtuin family of NAD^+^-dependent deacetylases (also referred to as the class III HDACs) have recently been demonstrated to remove a variety of noncanonical histone acyl PTMs, especially Sirtuin 2 and Sirtuin 3.^23,25,42^ Jennings et al. investigated the ability of the Sirtuins to remove the Kla modification from a non-histone peptide substrate and identified Sirtuin 2 (Sirt2) as a delactylase.^42^ Further research by Zu et al. revealed that Sirt2 also delactylates histones and that Sirt2 knockdown was sufficient to cause increased cellular accumulation of histone Kla in a neuroblastoma model.^25^ More recently, Du et al. demonstrated that Sirtuins 1 and 3 are capable of catalyzing lysine delactylation in histone and non-histone contexts.^43^ None of these studies identified the nuclear-localized members of the Sirtuin family, Sirtuins 6 and 7 (Sirt6 and Sirt7), as delactylases, but those that tested Sirt6 and Sirt7 all used peptide or single-histone substrates in their screens of deacylase enzymes. Experimental evidence indicates that Sirt6 exhibits very low deacylase activity on peptide and histone substrates but much higher deacylase activity on whole nucleosome substrates and that this behavior is dependent on Sirt6-histone and Sirt6-DNA interactions.^44–49^ Several recent structural studies further support those biochemical studies.^50–52^ These findings led us to hypothesize that Sirt6 and/or Sirt7 may exhibit previously overlooked delactylase activity when tested in a nucleosomal context. Herein, we performed biochemical and cell-based assays that show that Sirt6 is a histone delactylase.

## Results

### Sirt6 exhibits NAD^+^-dependent delactylase activity *in vitro*

We first tested whether Sirt6 is capable of delactylating histones *in vitro*. To accomplish this, we isolated nucleosomes from HEK-293T cells that were treated with 25 mM sodium L-lactate (to stimulate Kla) via detergent-free lysis, sonication, and digestion with micrococcal nuclease.^53^ These nucleosomes were then incubated with recombinantly expressed Sirt6 (full-length, human sequence), NAD^+^, or both, and histone Kla levels were measured using a pan-α-Kla antibody (**Figure 1A,B**). We observed that histone Kla levels were decreased only in the presence of Sirt6 and NAD^+^, leading us to conclude that the observed delactylation was due to the NAD^+^-dependent deacylase activity of the recombinant Sirt6.

**Figure 1.**
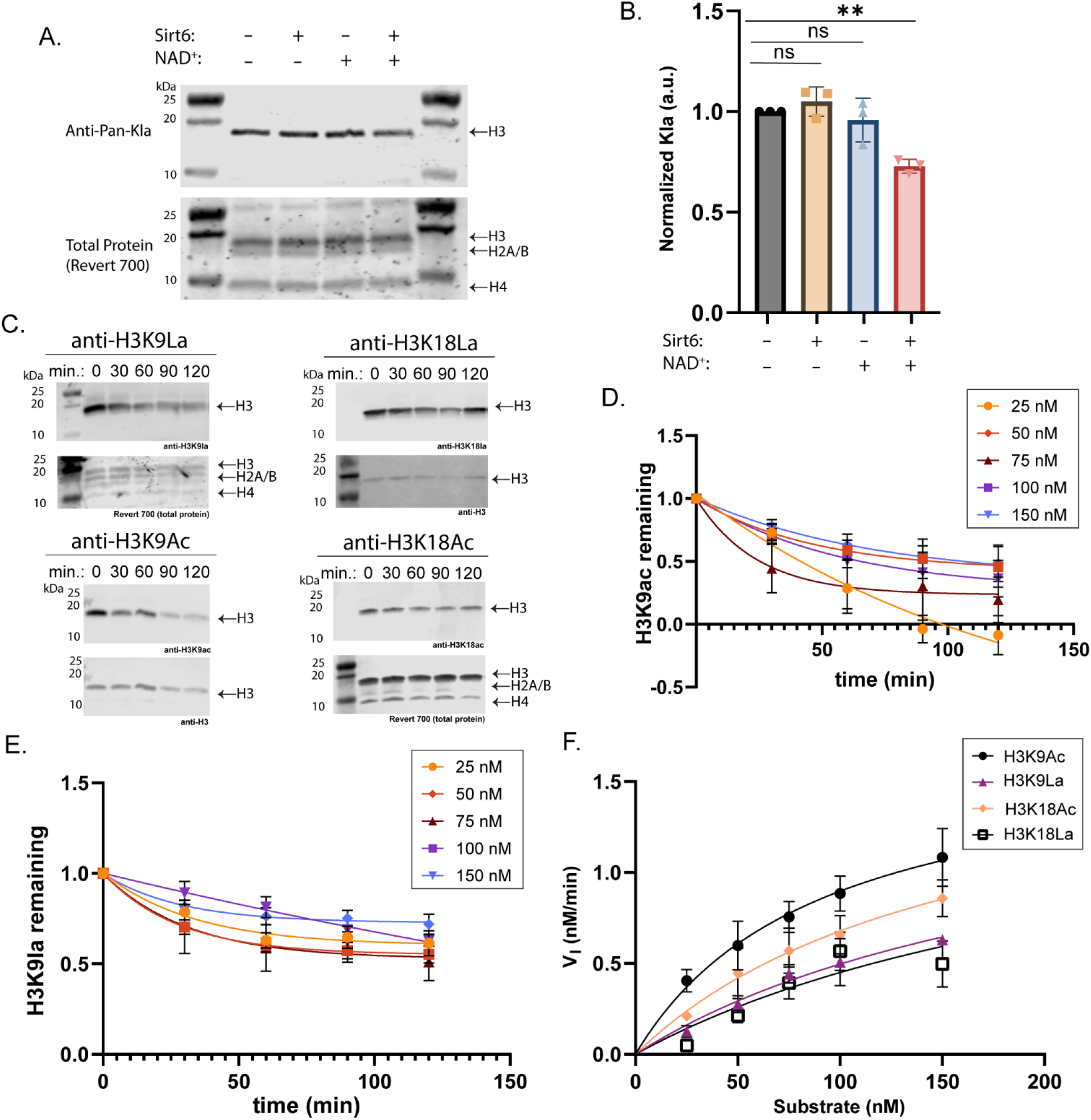
Sirt6 exhibits histone delactylase activity *in vitro*. (A) Western blot of histone Kla levels on nucleosomes isolated from HEK-293T cells that were incubated with recombinant Sirt6 and/or NAD^+^. (B) Quantitation of the blot from panel A. Densitometry data were corrected based on the total protein stain then normalized to the untreated control condition. Groups were compared using a one-way ANOVA followed by Tukey’s *post hoc* test. n=3, error plotted as S.D. p (-/-v. +/+) = 0.006. (C) Western blot analysis of time courses of enzymatic deacylation reactions using 25 nM substrate nucleosome, 100 nM Sirt6, and 1 mM NAD^+^. (D) Quantitation of western blot data for the H3K9Ac substrate in panel C. V_I_ was calculated for each substrate concentration by fitting the first three data points from this curve using standard linear regression. (E) As in panel D, but for the H3K9La substrate. (F) V_I_ values plotted as a function of substrate concentration for each PTM substrate. For panels D, E, and F, n=3, error plotted as S.D.

To assess which histone sites are delactylated by Sirt6, we incubated HEK-293T nucleosomes with recombinant Sirt6 and NAD^+^ and analyzed the samples using a panel of site-specific Kla and Kac antibodies (**Figure S1**).^21^ We were unable to detect several PTMs from the panel using these HEK-293T nucleosomes, so we also isolated nucleosomes from Sirt6-/-U2OS cells (“S6KO”) using the same protocol as for the HEK-293T nucleosomes. We detected several of the ‘missing’ PTMs in this cell line, leading us to conclude that those PTM levels were below the limit of detection in the HEK-293T cells. Other PTMs that were detected using HEK-293T nucleosomes were below the limit of detection on the U2OS nucleosomes. We found that Sirt6 deacetylates and delactylates at H3K9 and H3K18 but did not detect delactylation of H3K56 or H4K12 in the assay. For the PTMs at H3K9, which were detected in both cell lines, the activity of Sirt6 was consistent on nucleosomes derived from the different cell lines.

### Sirt6 is a better deacetylase than delactylase *in vitro*

To quantify the ability of Sirt6 to delactylate histones relative to its canonical deacetylase activities, we performed *in vitro* kinetics assays similar to previously published protocols.^50^ We prepared a panel of singly-modified mononucleosomes with H3K9ac, H3K9la, H3K18ac, or H3K18la by semisynthesis. These nucleosomes were incubated with recombinant Sirt6 (human, full-length) and NAD^+^ across a 2-hour time course at varying concentrations of nucleosome, and the extent of PTM removal was quantified using Western blot (**Figure 1C-E and S2A-B**). The time t=0 data were used to determine whether the antibodies used exhibited linear response across the substrate concentrations used in the kinetics assay (**Figure S2C**). The linear regression formula generated from the linear range testing was then treated as a standard curve, and the intensities from the rest of the kinetics data, corrected based on a loading control, were interpolated in the curve to calculate the amount of substrate remaining at each time point. Since the rate of substrate consumption was approximately linear during the first hour of data collection for all four substrates, the first three data points for each substrate were fit using standard linear regression to calculate an initial rate of deacylation (VI, **Figure 1F and S2D**). These VI values were then plotted as a function of substrate concentration and fit using the Michaelis-Menten model to determine an apparent KM,app and kcat,app for each substrate (**Table S1**). Based on this data, we determined that Sirt6 deacetylase activity is more efficient than its delactylase activity, and that this is primarily due to poorer binding to the Kla substrates. We also expressed human, full-length Sirt7 and performed similar assays profiling Sirt7 deacylase activity. We observed that Sirt7 exhibits deacetylase but not delactylase behavior at H3K18 (**Figure S2E-G**). As such, we did not proceed with kinetics analysis of Sirt7.

### Sirt6 deletion increases cellular accumulation of histone Kla

To assess the histone delactylase activity of Sirt6 and Sirt7 in a cell culture model, we knocked out Sirt6(- /-) or Sirt7(-/-) in U2OS cells (“S6KO” or “S7KO”). These cell lines were then treated with varying concentrations of sodium L-lactate for 24 h to stimulate histone Kla. Kla levels on acid-extracted histones from the cells were measured via immunoblotting and compared to the wild type U2OS cells (**Figure 2A,B**). The S6KO cells exhibited a greater accumulation of histone Kla compared to the WT or S7KO cells upon sodium L-lactate supplementation (**Figure 2C**). We also measured levels of histone Kac on the extracted histones (**Figure 2D**). The S6KO and S7KO cells exhibited no significant differences in total histone Kac compared to WT cells with or without lactate supplementation. This result supports the selectivity of the pan-Kla antibody and is consistent with previously published data from other cell lines treated with sodium L-lactate.^17^ It also suggests that this lactate supplementation paradigm leads to Kla increases at previously non-acetylated sites, as opposed to displacing existing Kac, at least at the bulk level. Even though the absence of Sirt6 led to higher accumulation of histone lactylation upon lactate supplementation, the S6KO cells did not exhibit a higher baseline level of total histone Kla without supplementation (**Figure 2E**). Since we did not observe evidence of Sirt7-catalyzed delactylation in either our *in vitro* or cell-based assays, subsequent experiments focused solely on the activity of Sirt6.

**Figure 2.**
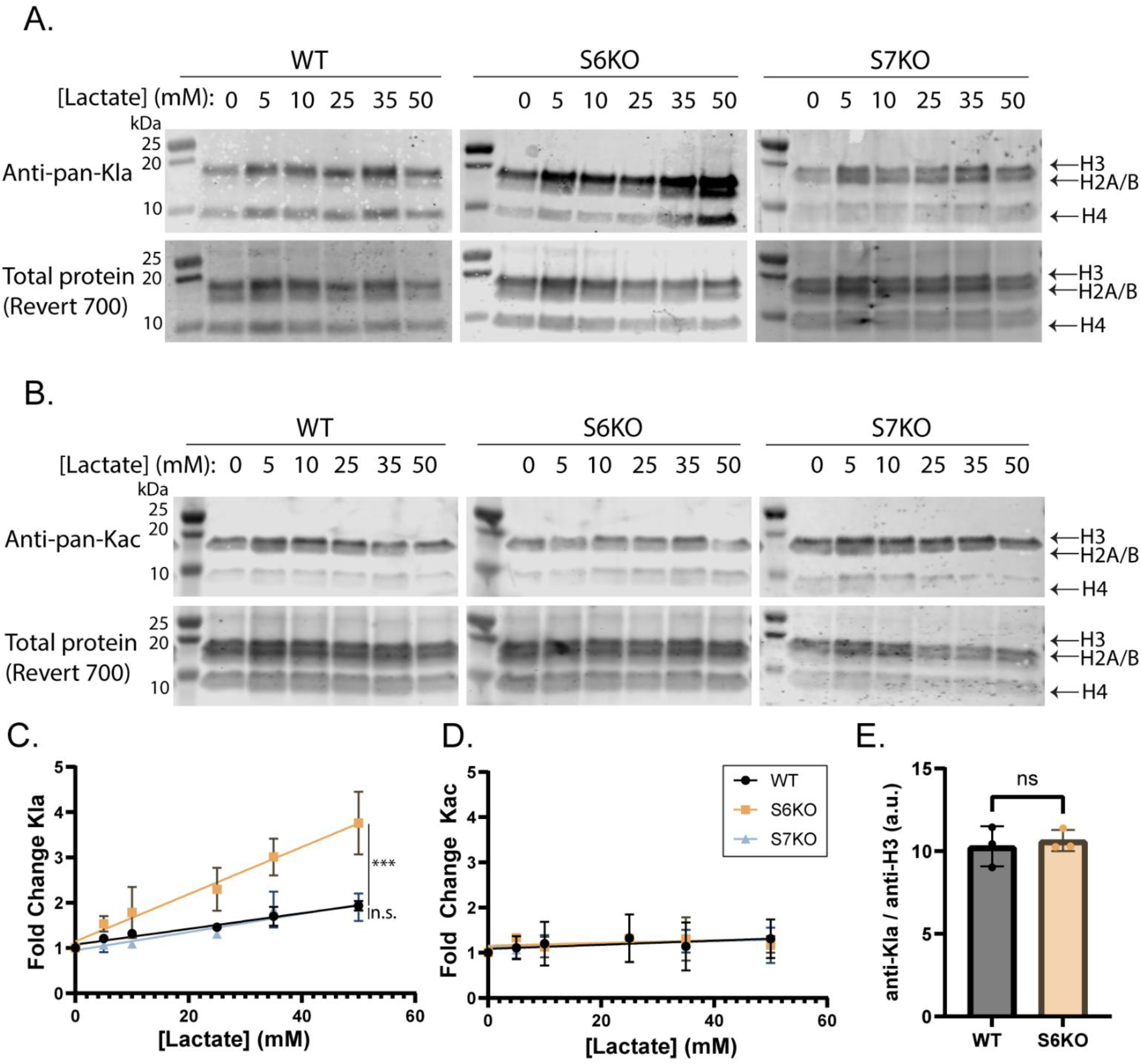
Sirt6 depletion leads to histone Kla accumulation. (A) Western blot using a pan-lactyllysine antibody to analyze Kla levels on acid-extracted histones from wild type (“WT”), Sirt6 knockout (“S6KO”), and Sirt7 knockout (“S7KO”) U2OS cells in the presence of a titration of sodium L-lactate. Total protein was measured using a fluorescent total protein stain. (B) As in panel A but using a pan-acetyllysine antibody to analyze levels of histone Kac. (C) Western blots from panel A quantified by normalizing Kla signal to total protein then represented as a fold-change from the untreated condition. n=3, error plotted as S.D. Slopes of linear regressions were compared using Welch’s t-test. p (slope, S6KO v. WT) < 0.0001. p (slope, S7KO v. WT) = 0.39. (D) Western blots from panel B quantified by normalizing Kac signal to total protein then represented as a fold-change from the untreated condition. For WT and S6KO, n=3. For S7KO, n=2. Error plotted as S.D. (E) Baseline histone lactylation in untreated cells was measured using a pan-Kla antibody as in panel A and normalized to the loading control. The blot images used to generate this plot are shown in Figure 5. Error plotted as S.D., p = 0.68.

Using the wild type and Sirt6-KO U2OS cells, we performed an experiment in which Kla was stimulated using rotenone, a mitochondrial Complex I inhibitor, to increase intracellular lactate as previously described.^17^ Under these conditions, we again observed that S6KO cells accumulated histone Kla, but not histone Kac, to a greater extent than WT cells (**Figure 3**). We also repeated the lactate supplementation experiment with the addition of sodium oxamate (a lactate dehydrogenase inhibitor). Under these conditions, the cells should not be able to produce lactate from metabolic pyruvate, and the supplemental lactate should not be able to be oxidized to pyruvate. Under these conditions, the S6KO cells again accumulated histone Kla to a higher extent than WT or S7KO cells (**Figure S3**).

**Figure 3.**
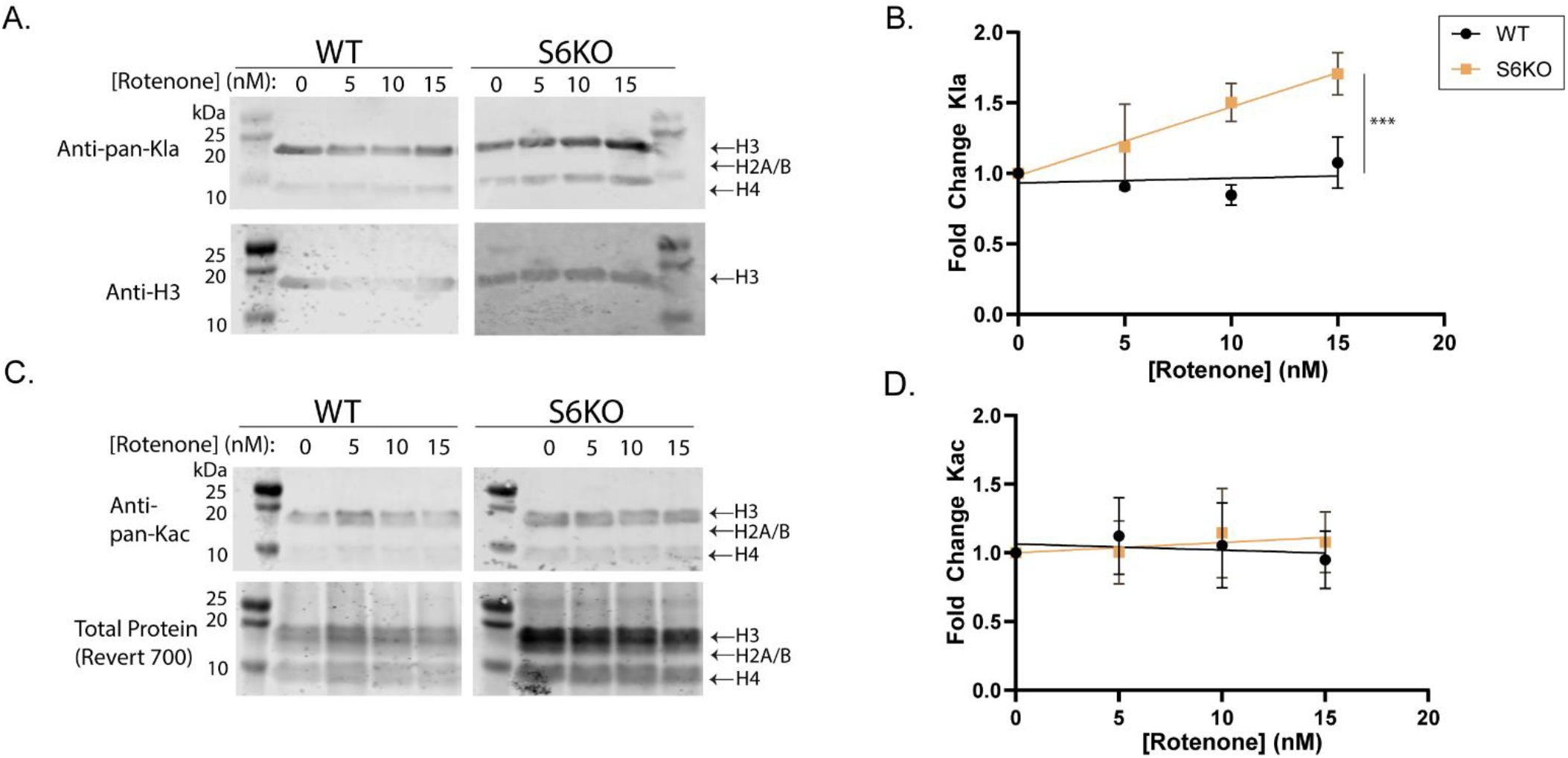
Sirt6 exhibits delactylase activity in cells. (A) Western blot using a pan-lactyllysine antibody (PTM BIO) to analyze Kla levels on acid-extracted histones from wild type and Sirt6 knockout U2OS cells in the presence of a titration of rotenone, a mitochondrial Complex I inhibitor. Total protein was measured using a fluorescent total protein stain (LI-COR). (B) Antibody signal was corrected based on the loading control signal, then normalized as a fold-change from the untreated condition. n=3, error plotted as S.D. Slopes of linear regressions were compared using Welch’s t-test, p=0.0005 (C) As in panel A but using the pan-acetyllysine antibody to measure levels of histone Kac. (D) Quantitation of data from panel C. Data were processed as in panel B. n=3, error plotted as S.D.

To determine the sites at which Sirt6 delactylates in cells, we measured the amount of acetylation and lactylation at various histone sites in the U2OS WT and S6KO cells using a panel of site-specific antibodies (PTM BIO, **Figure 4**). As with the pan-acetyl antibody (Figure 2C,D), we observed no changes in the acetyl sites (H3K9ac and H3K18ac) under the +/-lactate or +/-Sirt6 conditions. We observed accumulation of Kla at H3K9 and H3K18 in the cells under lactate supplementation that was enhanced in the S6KO cells compared to the WT cells, agreeing with the pan-lactyl antibody data (**Figure 2A,B**). These data indicate that Sirt6 can remove Kla from H3K9 and H3K18, in line with our *in vitro* data. With the H4K12la antibody, we detected no changes in +/-lactate or +/-Sirt6. We were unable to detect H3K56la in these cells even under lactate supplementation (data available as uncropped blots in **Supporting Information**).

**Figure 4.**
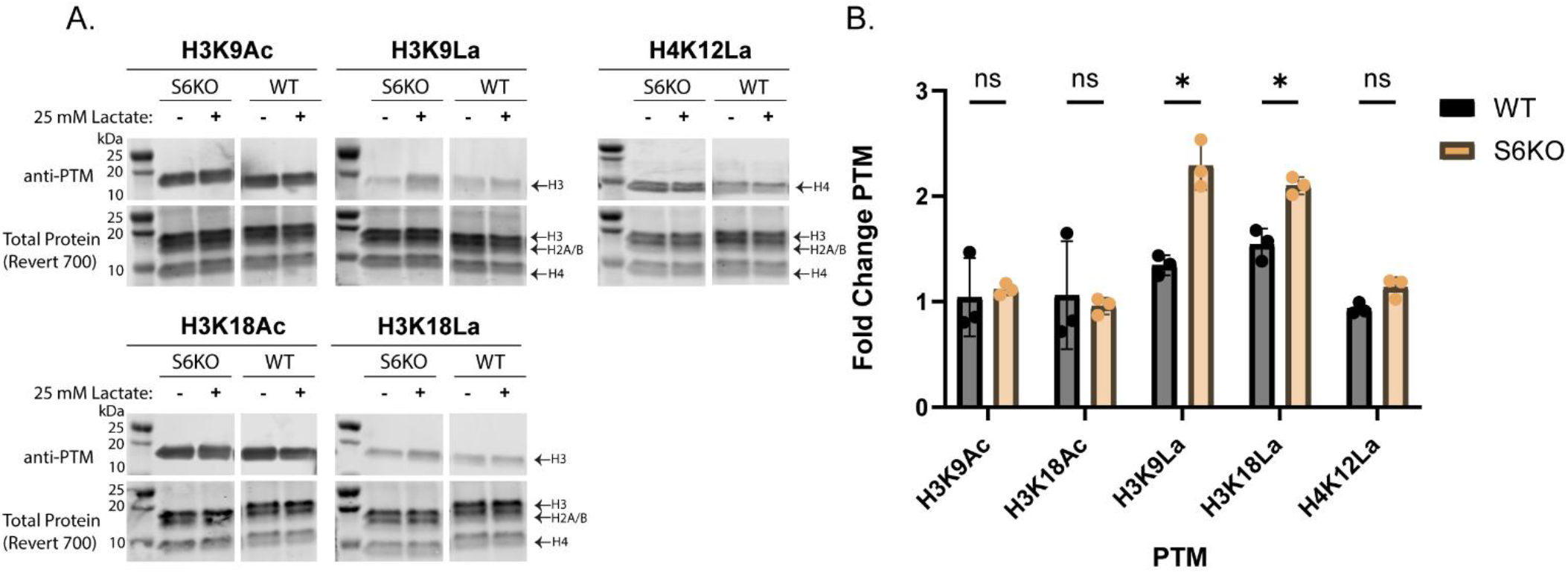
Sites of Sirt6-catalyzed deacylation in cells. (A) Levels of various histone acyl PTMs on acid-extracted histones from wild type or Sirt6 knockout U2OS cells were measured using site-specific antibodies (CST and PTM BIO). Total protein in each sample was measured using a fluorescent total protein stain (LI-COR). (B) Quantitation of data from panel A. Data were corrected based on the Revert 700 total protein stain, then normalized to the - Lactate control condition. Statistical analysis was performed using Student’s t-test and corrected for multiple hypothesis testing using the Holm-Šídák correction. n=3, error plotted as S.D. H3K9La p=0.013, H3K18La p=0.019.

### Sirt6 overexpression reduces cellular levels of Kla

We next sought to determine whether Sirt6 overexpression causes cells to exhibit decreased levels of histone Kla. We overexpressed Sirt6 (human, WT; CMV promoter) in WT U2OS cells (“WT-CMV-S6”) and in the S6KO cells (“S6KO-CMV-S6”) (**Figure 5**). Histones were extracted from untreated cells and from cells treated with 25 mM sodium L-lactate for 24 h, then analyzed for histone Kla. Sirt6 levels in each cell line were measured to validate the overexpression and to quantify the amount of Sirt6. In both treated and untreated cells, histone Kla was decreased in the overexpression and addback conditions (**Figure 5A-C,F-H**), supporting the hypothesis that Sirt6 is acting as a histone delactylase in the U2OS cells. In untreated cells, bulk histone Kac was also affected by changes in the amount of Sirt6 present in cells, though to a much lesser extent than Kla (**Figure 5D,E**). In cells treated with 25 mM lactate, the changes in Kla due to Sirt6 were much more pronounced. In these cells, histone Kla was inversely proportional to the amount of Sirt6 expression in the cell (**Figure 5I**).

**Figure 5.**
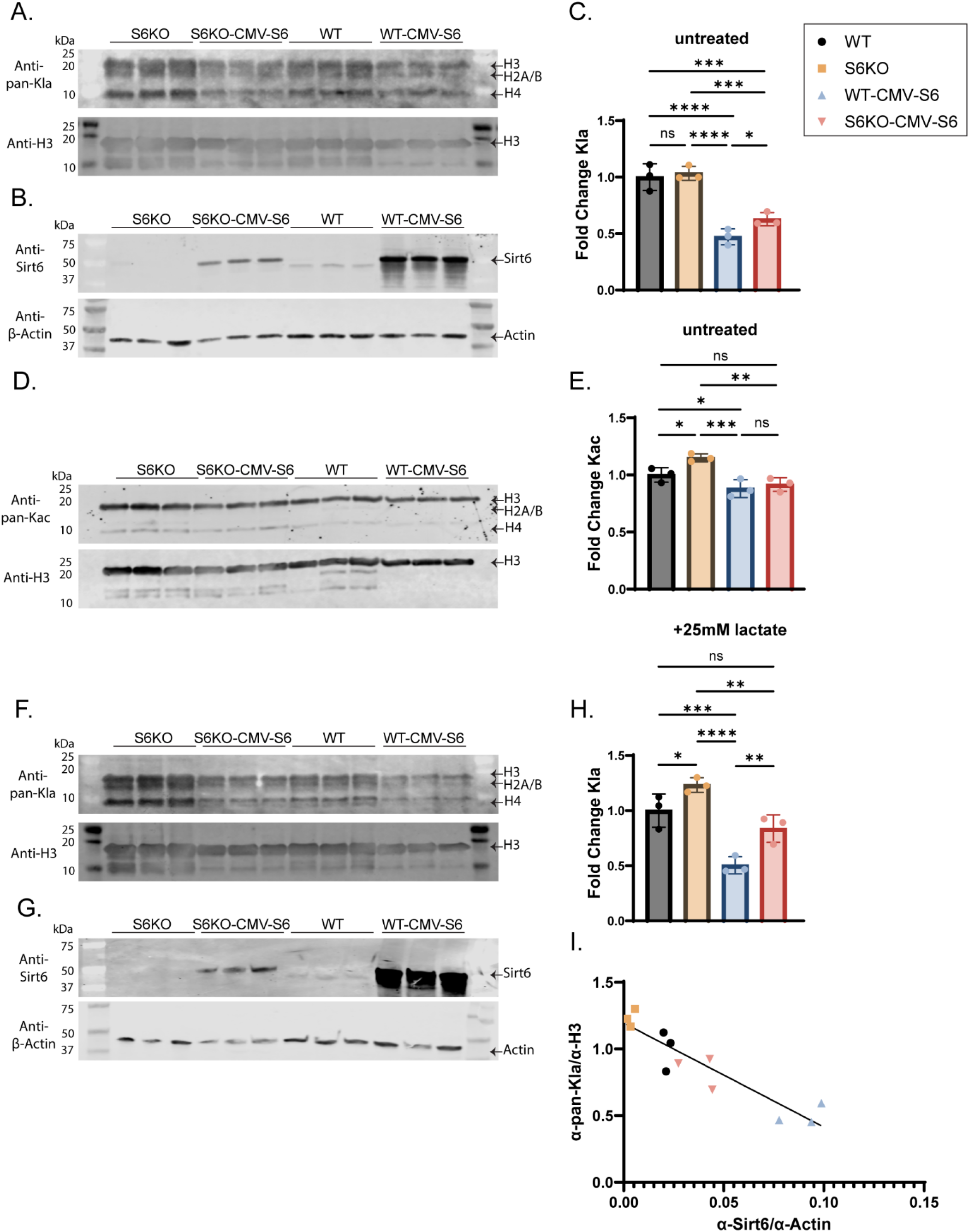
Cellular histone Kla is reduced by overexpression of Sirt6. (A) Western blot measuring histone Kla on acid-extracted histones in WT and Sirt6-KO U2OS cell lines not treated with supplemental L-lactate with and without overexpression of human Sir6 under the control of a CMV promoter. WT histone H3 (CST) is used as a loading control. (B) Western blot measuring Sirt6 in the soluble protein fraction from cell lines from (a). β-Actin is used as a loading control. (C) Kla signal from (a) was quantified, normalized to the loading control, and represented as a fold change from the average of the WT condition. Statistical analysis was performed using one-way ANOVA followed by Tukey’s *post hoc* test. (D) as in (a) but measuring histone Kac. (E) as in (c), but for Kac data from (d) (F) As in (a), but the cells were treated with 25 mM sodium L-lactate for 24 hours prior to histone extraction. (G) As in (b), but the cells were treated with 25 mM sodium L-lactate for 24 hours prior to cell lysis. (H) Kla signal from (f) was quantified, normalized to the loading control, and represented as a fold change from the average of the WT condition. (I) Inverse correlation of data from (f) and (g). Sirt6 signal was normalized by dividing by the β-actin loading control. Kla signal was normalized as in (h). The data were analyzed using a standard linear regression, R_2_ = 0.84.

### Sirt6 and Class I HDACs exhibit additive delactylase activity

The Kla and Kac data from the cell lines not treated with lactate (**Figure 5C**,**E**) seemed to disagree with the kinetics data (**Figure 1F**) that indicated Sirt6 is a better deacetylase than delactylase. These results led us to hypothesize that the U2OS cells are more effective at compensating for the loss of Sirt6 deacetylase activity, but that there is a lower capacity to compensate for loss of Sirt6 delactylase activity. We tested this hypothesis by investigating the ability of different enzymes to delactylate nucleosomes in mammalian cells. We treated the WT and S6KO U2OS cells with sodium L-lactate and with Panobinostat, a nonselective inhibitor of the Zn^2+^-dependent HDACs (Classes I, II, IV) and again measured global histone acetylation and lactylation levels (**Figure 6A-D**). Unsurprisingly, panobinostat increased the accumulation of histone Kac and Kla in both cell lines, and we observed that the Sirt6 knockout did not significantly affect histone acetylation levels (**Figure 6D and S4**), in agreement with the prior experiments. However, even in the presence of panobinostat, Sirt6 knockout still resulted in greater accumulation of histone Kla (**Figure 6C and S4**). This result indicates that the Zn^2+^-dependent HDACs cannot fully compensate for the loss of Sirt6 histone delactylation activity. It suggests that Sirt6 and the Zn^2+^-dependent HDACs may target different sites of lactylation within histones and/or loci within the genome. We also analyzed PTM levels on the selection of histone sites from **Figure 4** under Panobinostat treatment (**Figure S4**). We observed that at H3K9 and H3K18, the trends observed with the site-specific antibodies mirror the trend observed using the pan-PTM antibodies.

**Figure 6.**
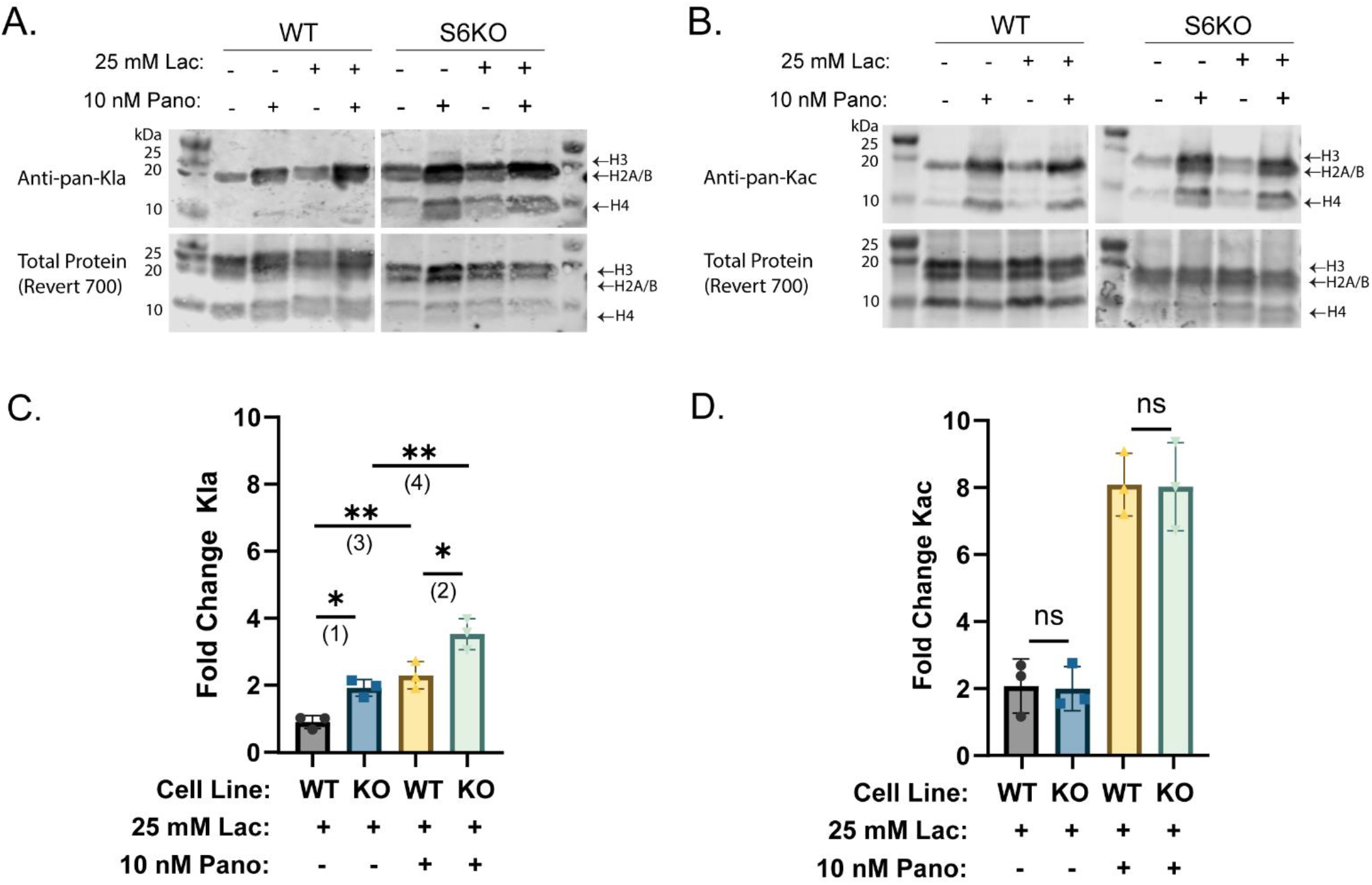
Sirt6- and HDAC-catalyzed delactylation are additive. (A) Western blot measuring histone Kla in WT and Sirt6-KO U2OS cells treated with sodium L-lactate and panobinostat, as indicated. A total protein stain (LI-COR) was used as a loading control. (B) As in panel A but measuring histone Kac. (C) Quantitation of selected data from panel A. Histone Kla signal was quantified using densitometry and corrected based on the total protein stain. Data are presented as a fold change from the untreated (-lactate, -panobinostat) condition. Statistical analysis was performed using one-way ANOVA followed by Tukey’s *post hoc* test. p_1_ = 0.03, p_2_ = 0.02, p_3_=0.005, p_4_=0.002. n=3, error plotted as s.d. (D) Quantitation of selected data from panel B. Data was processed as in panel C. n=3, error plotted as s.d. Quantitation of additional conditions is available in figure S4.

## Discussion

Previous studies seemed to ‘rule out’ Sirt6 as a histone delactylase based on *in vitro* assays. However, these assays either used non-histone peptide substrates^42^ or relied on using nicotinamide as a nonselective sirtuin inhibitor.^41^ Biochemical data indicated the existence of Sirt6-DNA and Sirt6-acidic patch interactions that are important for binding of Sirt6 to substrates and demonstrated that Sirt6 is much more active on nucleosome substrates than histone peptides,^44,45,47^ and recent cryo-electron microscopy studies confirmed these biochemical data.^50,51^ Using nucleosome substrates, we detected and characterized the delactylase activity of Sirt6. We determined the kinetic parameters of Sirt6 for various PTM substrates, finding that Sirt6 has somewhat poorer binding to lactylated nucleosomes than acetylated nucleosomes *in vitro*. Additionally, our data indicated that Sirt6 delactylates nucleosomes in cell culture at the H3K9 and H3K18 positions and that this delactylase behavior is additive with the delactylase behavior of the Zn^2+^-dependent HDACs. Many of our cell culture experiments were carried out with the addition of 25 mM sodium L-lactate to the media. While this is well above the normal human serum level of lactate (1.5-3 mM), lactate concentrations from 10-50 mM are known to occur in inflammation and cancers,^54,55^ and many of the phenotypes and diseases in which histone lactylation have been studied are closely connected to inflammation or cancer development and progression. Given this information, the lactate supplementation paradigm is a useful model to study effects of or regulators of histone lactylation.

Sirt6 has several experimentally demonstrated enzymatic functions including deacetylation, fatty deacylation, and ADP-ribosylation.^56–59^ These enzymatic activities ultimately affect a variety of biological processes. Sirt6 is known to regulate metabolic homeostasis, and it has been implicated in regulating DNA repair processes and metabolic processes such as insulin secretion and ketogenesis.^58,60–65^ Sirt6 is specifically known to downregulate glycolytic activity in cells by downregulating the activity of Hif1α.^66,67^ Given its role in regulating the expression of metabolic enzymes, it is plausible that Sirt6-catalyzed delactylation is part of a mechanism by which Sirt6 may regulate metabolic genes. Our data from cells not treated with sodium lactate indicate that Sirt6 ablation affects histone lactylation to a greater extent than histone acetylation in the U2OS cells, making it a strong potential candidate to target to selectively modulate levels of the histone Kla PTM. Manipulation of Sirt6 activity is highly experimentally tractable by genetic manipulation or by selective pharmacological inhibition.^68–70^ While it is unlikely that Sirt6 directly affects all instances of epigenetic regulation via histone Kla, it may serve as a useful control to deconvolute the effects of histone Kac from histone Kla in future studies, especially at the H3K9 and H3K18 positions.

## Materials and Methods

### Materials

All salt, buffers, and other chemicals were obtained from Fisher Scientific unless otherwise noted. All reagents used for solid-phase peptide synthesis (SPPS) were obtained from ChemImpex International unless otherwise noted. Other reagents for biochemical assays, cloning, and protein purification are listed in **Table 1**. Antibodies (with dilutions used) are listed in **Table 2**. The Sirt6 plasmid for *E. coli* expression was described in a previous study.^71^ SIRT7.4 was a gift from John Denu (Addgene plasmid # 13740 ; http://n2t.net/addgene:13740 ; RRID:Addgene_13740).^72^ The histone plasmids were described in a previous study.^73^ pLJM1 was a gift from David Sabatini (Addgene plasmid # 19319).^74^ pSpCas9(BB)-2A-Puro (PX459) V2.0 was a gift from Feng Zhang (Addgene plasmid # 62988 ; http://n2t.net/addgene:62988 ; RRID:Addgene_62988).^75^ The HEK-293T cells were a gift from Jared Rutter. The U2OS cells were a gift from Glen Liszczak.

**Table 1.**
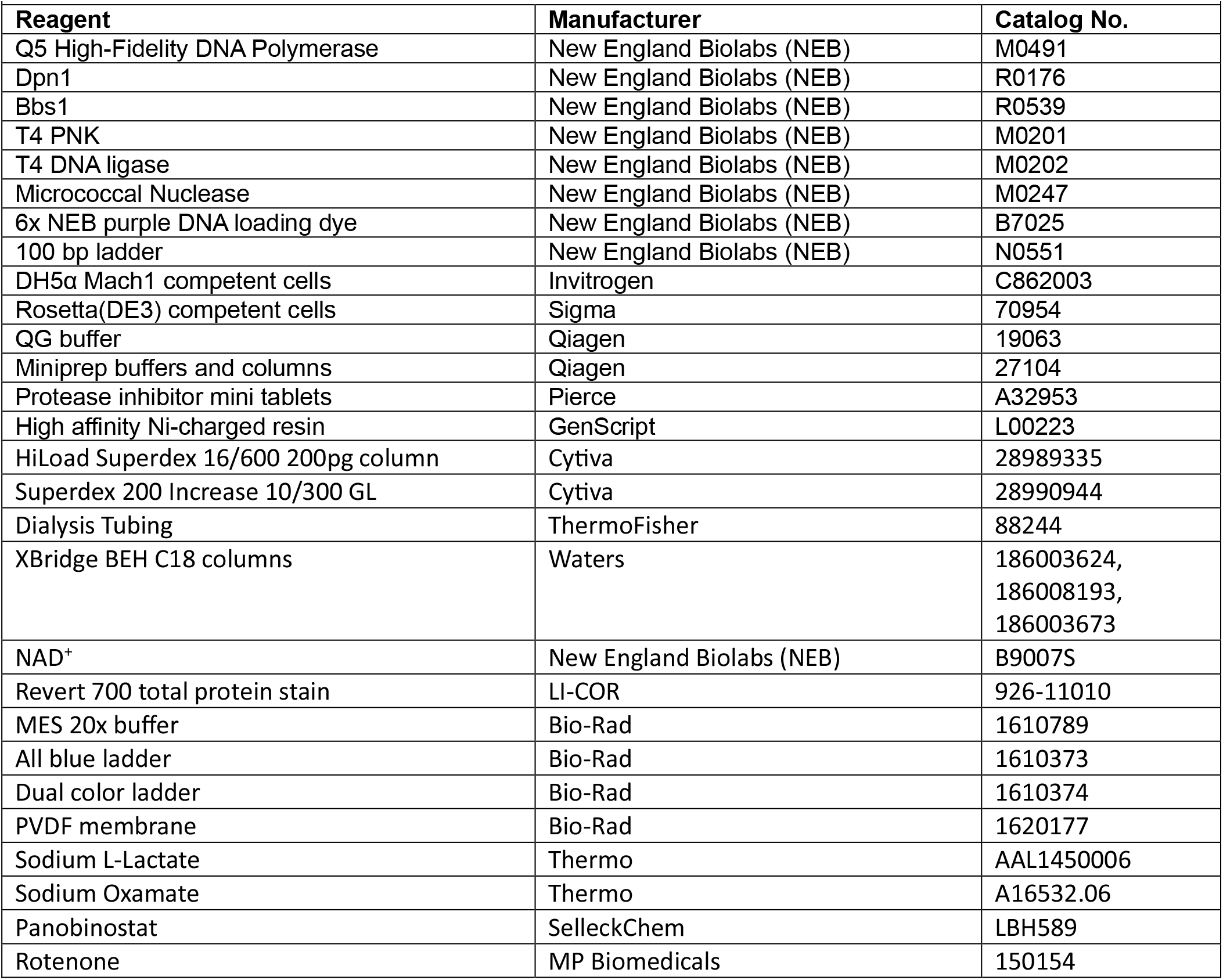
Materials

**Table 2.**
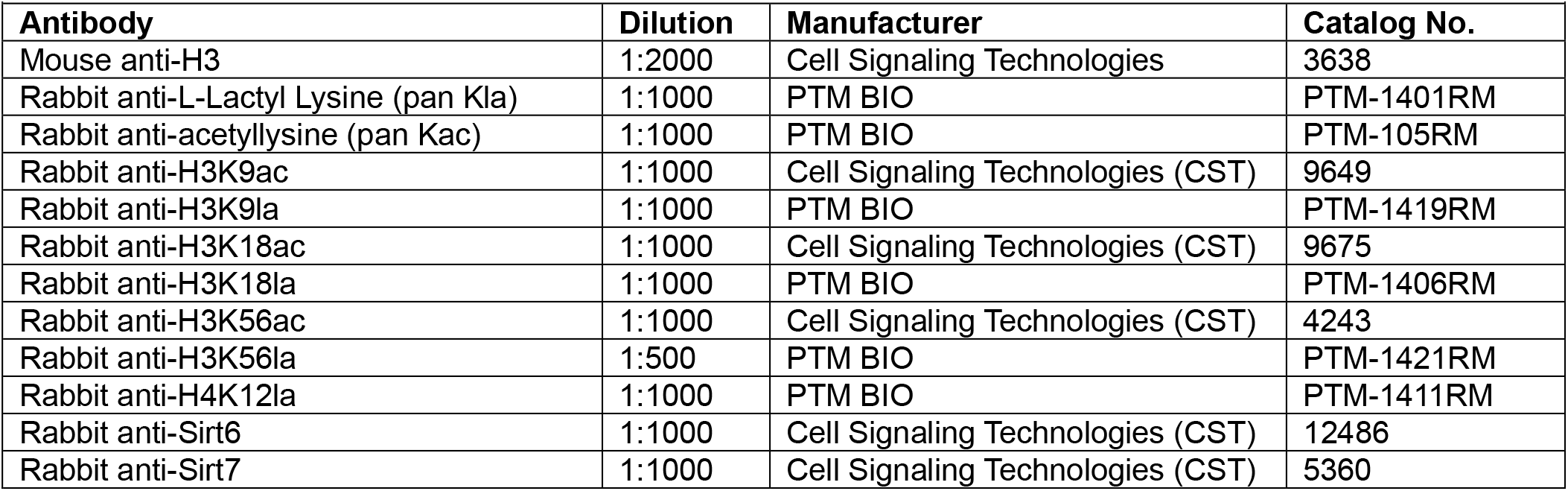
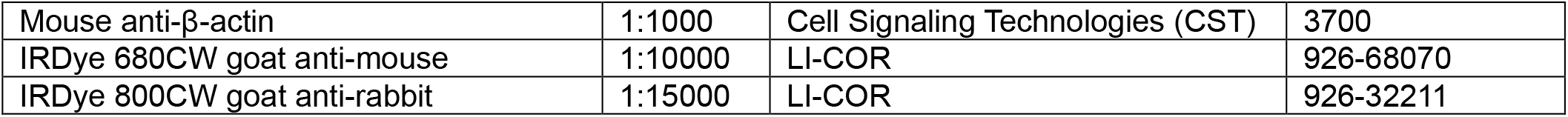
Antibodies

### Methods

#### Cloning

Q5 High-Fidelity DNA Polymerase (NEB) was used for all PCR amplification steps. All primers were obtained from the DNA/Peptide Core at the University of Utah. PCR products were treated with Dpn1 (NEB), and linear products were purified via a 1% agarose gel, extracted with QG buffer (Qiagen), and purified by spin column (Qiagen). NEBuilder HiFi DNA Assembly Master Mix (NEB) was used as described by the manufacturer. For blunt-end ligations, T4 PNK (NEB) and T4 DNA ligase (NEB) were used as indicated by the manufacturer. Reactions were transformed into DH5α Mach1 *E. coli* cells (Invitrogen) and plated on antibiotic-containing agar to select colonies for sequence verification. Plasmids were isolated from liquid cultures using Qiagen Miniprep buffers and spin columns as directed by the manufacturer. All plasmids were sequence verified by GENEWIZ (Aventa Life Sciences). Amino acid sequences are listed in the **Supporting Information**.

The pET28a-LIC-6x-His-SUMO-Sirt6 was cloned as described previously. The pQE-80-6xHis-Sirt7(Δexon2) was Addgene plasmid #13740. The missing exon 2 was synthesized as a GeneBlock (IDT) and cloned into the Sirt7 gene. A SUMO tag was also added between the 6xHis and the Sirt7 to yield pQE-80-6xHis-SUMO-Sirt7. For mammalian cell overexpression, the full length, human Sirt6 sequence with a C-terminal FLAG tag was cloned into a pLJM1 mammalian expression vector with a CMV promoter (“pLJM1-Sirt6-3xFLAG”).,

For the human cell knockouts, the following sgRNA sequences were cloned into the pSpCas9(BB)-2A-Puro (PX459) V2.0 vector:

Sirt6: sgRNA1-CCTGAAGTCGGGGATGCCAG,^76^ sgRNA2-TACGTCCGAGACACAGTCGT^77^

Sirt7: sgRNA1-CGTTACCAGGTCCGCGCTCT, sgRNA2-GCTTCAGGCCCTCGCGCCGC, sgRNA3-GGCCCTGCAGCTCCGTTACC^78^

The oligonucleotide design and cloning were performed according to the protocol provided by the Zhang lab on the Addgene #62988 website. Briefly, the oligonucleotides were annealed and phosphorylated using T4 PNK (NEB) following manufacturer’s protocol. 1 µg of the PX330 Cas9 plasmid was digested with *Bbs*I (NEB) following manufacturer’s protocol and purified by agarose gel. Digested plasmid and phosphorylated/annealed oligo inserts were ligated using the T4 DNA Ligase (NEB) according to manufacturer’s directions. The resulting plasmids were transformed into chemically competent DH5α *E. coli* cells. Colonies were selected and plasmid identities were sequenced for verification.

### Recombinant protein expression

Plasmids were transformed into BL21 Rosetta (DE3) *E. coli* cells (Millipore) and grown to OD 0.6 in LB Miller (Fisher), shaking at 180 rpm at 37°C with the addition of the appropriate antibiotic for selection. The temperature was then lowered to 18°C, and cells were induced by addition of 0.5 mM IPTG (Fisher) and allowed to grow for 18 hours before collection and storage at -80°C.

### Protein purifications

#### Histones

Full length, human histones H2A, H2B, H3, and H4, as well as H3[ΔN, 1-14]A15C and H3[ΔN, 1-28]A29C were purified as described previously.^73,79^

#### Sirtuins

Full length, human Sirt6 was expressed and purified as previously described. Briefly, cells containing the expressed 6xHis-SUMO-Sirt6 were lysed in lysis buffer (50 mM Tris, pH 7.5, 500 mM NaCl, 10 mM MgCl_2_, 5mM BME, 1 mM PMSF). Sirt6 was purified using a nickel affinity resin (Genscript) and eluted in lysis buffer with 300 mM imidazole added. Ulp1 was added to the elution to cleave the 6x-His-SUMO tag, then the solution was dialyzed against lysis buffer with no imidazole or PMSF. The cleaved 6x-His-SUMO was removed using a reverse nickel affinity purification. The purified, cleaved Sirt6 was concentrated and analyzed on a HiLoad 16/600 Superdex 200 pg size exclusion column (Cytiva) and eluted in storage buffer (50 mM Tris, pH 7.5, 100 mM NaCl, 1 mM MgCl_2_., 10% glycerol, 2 mM DTT). Fractions containing Sirt6 were pooled, concentrated, and stored at -80°C. Sirt7 was purified using the same protocol, except lysis and storage buffer did not contain MgCl_2_. A truncation product was detected, which was not able to be removed using SEC despite its apparently small size, but this did not seem to impact the activity of the isolated Sirt7. Purified proteins were analyzed via SDS-PAGE (**Figure S5A**).

### Histone H3 semisynthesis

The following histone H3 peptides were synthesized using a standard Fmoc-protected SPPS strategy as previously described.^73,79^ The peptides were synthesized on a hydroxytrityl resin (Chemmatrix) functionalized with hydrazine.

H3(1-14, K9alloc): ARTKQTARK(alloc)STGGK-NHNH_2_.

H3(1-28, K18alloc): ARTKQTARKSTGGKAPRK(alloc)QLATKAARKS-NHNH2.

After synthesis, the alloc protecting group was removed from the peptides on resin using palladium tetrakistriphenylphospine and dimethylbarbituric acid as previously described.^73,79^ The K9ac and K18ac peptides were generated by coupling acetate to the deprotected ε-amine using 10% Acetic Anhydride/20% DIPEA in DMF. The K9lac and K18lac peptides were generated by coupling L-lactate to the deprotected ε-amine using 6 eq L-lactic acid, 5.5 eq PyAOP, and 12 eq of DIPEA in DMF. The peptides were then cleaved and fully deprotected using trifluoroacetic acid (95% TFA, 2.5% water, 2.5% triisopropylsilane).

The resulting hydrazide peptides were converted to thioester peptides following previously published protocols.^73,79^ Briefly, each peptide was oxidized using NaNO_2_, then reacted with sodium mercaptoethane sulfonate (MESNa) to generate a thioester-containing peptide for native chemical ligation. The thioester peptides were purified using RP-HPLC (Agilent 1260) and characterized via LC-MS (**Figure S6**). The MESNa peptides were then reacted with truncated histone proteins (H3[ΔN, 1-14]A15C for the H3K9ac and H3K9la peptides, H3[ΔN, 1-28]A29C for the H3K18ac and H3K18la peptides) following previously published protocols for native chemical ligation. Briefly, each peptide was reacted with the corresponding truncated histone and trifluoroethanethiol to generate the full-length histone, then the histones were subjected to radical desulfurization to yield the native H3A15 or H3A29. The resulting histones were purified via RP-HPLC (Agilent 1260 Infinity II) and characterized using LC-MS (Waters Acquity QDa and Waters Xevo G2-XS QTof) (**Figure S7**).

#### Octamer/nucleosome formation

Wild type, H3K9ac, H3K9la, HK18ac, and K3K18la octamers were assembled by combining H3 (WT or modified) with H2A, H2B, and H4 and dialyzing as previously described.^73,79^ Octamers were purified by size exclusion chromatography (Superdex 200 increase column, Cytiva). Mononucleosomes were assembled by adding ds601 DNA (sequence in **Supporting Information**) and dialyzing as previously described. The mononucleosomes were analyzed on a native 5% TBE gel to assess their quality (**Figure S5C**).

#### Tissue Culture

U2OS and HEK 293T cell lines were grown in Dulbecco’s modified Eagle’s media (DMEM, Gibco) supplemented with 10% fetal bovine serum (Gibco) and 1% penicillin/streptomycin (Gibco). Cells were routinely tested for mycoplasma contamination and periodically genotyped using STR analysis (ATCC). Cells were subcultured at 95% confluence; U2OS cells were routinely subcultured using a 1:8 dilution and HEK-293T cells were routinely subcultured using a 1:10 dilution. For all treatment experiments, cells were plated at least 24 h prior to treatment and allowed to grow to 80% confluence. Cells were subcultured for ≤ 12 passages. Cells treated with sodium L-lactate, sodium oxamate, or Panobinostat were treated for 24 h immediately prior to harvesting. Cells treated with rotenone were treated for 4 h immediately prior to harvesting. Control cells were treated with vehicle (1% sterile PBS for sodium lactate and sodium oxamate, 0.5% DMSO for Panobinostat and rotenone). All cell experiments were performed with three biological replicates. Cell lysis and histone acid extraction were performed using the EpiQuik total histone extraction kit (EpiGentek) according to manufacturer’s published protocols. Soluble lysates were obtained by preserving the supernatant from the initial lysis step of the histone extraction procedure (“Pre-lysis” according to the manufacturer’s nomenclature).

#### Nucleosome isolation

Approximately 3.0 × 10^7^ cells were lysed in hypotonic buffer (10 mM Tris, pH 7.5, 10 mM KCl, 1.5 mM MgCl_2_, 0.34 M sucrose, 10% glycerol, 1 mM DTT, 1x Pierce mini protease inhibitor tablet) with 0.1% NP-40 added.^53^ Cells were left on ice for 10 min then pelleted at 1300 rcf for 5 min. The pellet was washed in hypotonic buffer, then resuspended in chromatin precipitation buffer (3 mM EDTA, 0.2 mM EGTA, 1 mM DTT, Pierce mini protease inhibitor tablet). The pellet was pelleted at 1700 rcf for 5 min, then washed with precipitation buffer again. The pellet was washed once with resuspension buffer (hypotonic buffer + 2 mM CaCl_2_). The suspension was sonicated with a microtip rod sonicator (40% amplitude, 15 s. on, 45 s. off, 6 cycles). Micrococcal nuclease was added (2000 u, 1 µL) and the mixture was incubated at 37°C for 30 min. The reaction was quenched by adding EGTA to 10 mM. The suspension was then pelleted at 17,000 rcf for 5 min, and the nucleosomes were collected in the supernatant. These nucleosomes were also analyzed on a native 5% TBE gel to assess their size and quality (**Figure S5C**)

### CRISPR-Cas9 knockouts

For the Sirt6 knockout, separate 10-cm plates of U2OS cells were transfected with each Cas9/sgRNA plasmid. For the Sirt7 knockout, one 10-cm plate was transfected with the pooled sgRNA plasmids. Cells were at 90% confluence at transfection. Cells were transfected using Lipofectamine 3000 (Invitrogen) in Optimem (Gibco) following the manufacturer’s protocol. Cells received 15 µg total DNA per 10-cm plate. Cells were left in Lipofectamine 3000/DNA/Optimem for 6 h, then recovered in antibiotic free complete DMEM. Beginning at 48 h post-transfection, the cells were selected using puromycin (1 µg/mL) in complete media.

For Sirt7 KO cells, the selection was continued for 96 h. The knockout was confirmed in the polyclonal cells by western blot (**Figure S5B**). For Sirt6 KO cells, after a 48-h selection, the cells were plated by dilution in 96-well plates to yield single colonies in DMEM without any puromycin. The single colonies were expanded and screened for Sirt6 KO by western blot. All experiments were performed from one clonal cell line (designated ‘A6’). Verification of Sirt6 knockout in this clone is in **Figure 5B**.

### Sirt6 overexpression

Wild-type and S6KO U2OS cells were plated in a 6-well plate at 2.0 × 10^5^ cells per well. Once cells reached 90% confluence, cells were transfected with the Sirt6 plasmid using Lipofectamine 3000 (Invitrogen) and Optimem (Gibco), following manufacturer’s recommended protocol. Each well of cells received 1 µg DNA. Cells were left in Optimem for 6 h, then recovered in antibiotic free complete DMEM overnight. Cells were then incubated in complete DMEM for 48 h and harvested. Control cells (i.e. cells not receiving the overexpression plasmid) were subjected to the same treatment with lipofectamine, but with no addition of DNA, and received all the same washes and media changes as the treated cells.

### *In vitro* sirtuin activity assays

For basic assays testing whether Sirt6 or Sirt7 is active on a substrate, stock solutions of recombinant sirtuin (30 µM, 10x), NAD^+^ (10 mM, 10x), and 5x reaction buffer (250 mM Tris, pH 7.5, 100 mM NaCl, 10 mM MgCl_2_, 10 mM DTT) were prepared. The reaction buffer and stock solutions were diluted in milliQ water, and substrate nucleosome was added to a final concentration of 100 nM (for semisynthetic nucleosomes) or 500 nM (for nucleosomes isolated from cells). The reactions were performed in 30 µL volumes, and were incubated at 37°C for 2 h, then quenched by addition of 10 µL 5x SDS loading buffer (250 mM Tris, pH 6.8, 10% w/v SDS, 30% v/v glycerol, bromophenol blue, 7.5% v/v BME). For kinetics experiments, the above assay was adjusted to be performed in a 96-well plate. A 5x master mix was prepared containing Sirt6 (500 nM), NAD^+^ (5 mM), and buffer components (250 mM Tris, pH 7.5, 100 mM NaCl, 10 mM MgCl_2_, 10 mM DTT). To 16 µL of the master mix was added substrate nucleosome to desired final concentration (25-150 nM) and milliQ water to reach 80 µL. Reaction times were calculated from the addition of substrate. Reactions were incubated at 37°C and quenched by addition of 20 µL 5x SDS loading buffer at indicated time points. 96-well plate assays were loaded with milliQ water in the outer wells adjacent to the edge of the plate to avoid uneven evaporation of water from reactions. All assays were performed with 3 technical replicates.

#### Western blotting

Samples were denatured by boiling in standard SDS loading buffer (50 mM Tris, pH 6.8, 2% w/v SDS, 6% v/v glycerol, bromophenol blue, 1.5% v/v BME) for 1 min. Samples were analyzed on either a 12% Bis Tris PAGE gel (for histone analytes) or a 4-12% Bis Tris PAGE gel (for all other analytes) using 1x MES running buffer (Bio-Rad) at 180 V. Samples were transferred to a PVDF membrane using semi-dry transfer at 25 V for 30 min, then blocked with 3% dry milk in TBST (50 mM Tris, pH 7.5, 150 mM NaCl, 0.05% Tween-20) for 1 h. Membranes were washed 3x for 5 min with TBST, then incubated with primary antibody at indicated dilution for 1-2 h at RT. Membranes were again washed 3x with TBST, then incubated with secondary antibody (1:10,000 for mouse secondary, 1:15,000 for rabbit secondary) for 1 h at RT. The membranes were again washed 2x with TBST and 1x with milliQ water for 5 min, then imaged using a LI-COR Odyssey imager. Cross-target reactivity profiles of site-specific antibodies from PTM Bio are available from the manufacturer.

### Software/Data analysis

Quantitation of western blots was performed using densitometry tools in the ImageStudio software package (LI-COR). All quantified western blots were corrected by dividing the analyte signal by a loading control (typically histone H3 or Revert 700 fluorescent total protein stain). Following the correction, the data was either presented as a corrected fluorescence value (in a.u.) when all data was acquired on the same blot or normalized as a fold change from a control condition to allow better comparison between different blots/antibodies. All densitometry quantitations and corrections performed are available as **Supplemental Data File 1**. All plotting and statistical analysis was performed using GraphPad Prism. Linear fits were performed using simple linear regression (unconstrained) in GraphPad Prism. Kinetics analyses were performed by first corrected for loading within each replicate by dividing the loading control densitometry value from each lane by the mean of the loading control values for that replicate, then multiplying the densitometry value of the PTM signal by that correction ratio. The densitometry values for the t=0 time points were then used to generate a standard curve for each antibody. The remaining data points were then converted to concentrations of substrate by interpolating within the standard curve. The calculated concentration values were then fit using simple linear regression on the first three data points (R^2^ ≥ 0.8 for all fits) to find V_I, app_. The V_I_ values were then fitted using the Michaelis-Menten kinetics function in Prism (unconstrained) to calculate apparent kinetics parameters. Statistical comparisons were performed using Student’s t test analysis when comparing two groups, Student’s t-test followed by multiple comparison correction when making multiple comparisons between the same two groups, and one-way ANOVA with an appropriate post hoc test (as noted in figure captions) for comparing more than two groups.

## Author Contributions

G.A.N. and K.L.D. designed the experiments. K.L.D. generated the Sirt6 knockout U2OS cell line. G.A.N. generated the Sirt7 knockout cell line and performed the cell culture experiments, *in vitro* enzyme assays, kinetics assays, and protein semisynthesis. F. and Z.Y. assisted in some of the protein semisynthesis and *in vitro* enzyme assays. N.P. performed the cloning for CMV-Sirt6 overexpression and the purification of recombinant Sirt6. N.P. and J.B. performed the cloning and purification of recombinant Sirt7. G.A.N. and K.L.D. prepared the manuscript with input from all authors.

## Acknowledgements

The authors acknowledge Dr. Bailey Miller and Dr. Eric Schmidt for assistance with mass spectrometric analysis of the histone peptides and proteins. We acknowledge the DNA/Peptide Synthesis Core at the University of Utah.

## Funding

National Institutes of Health grant R35GM143080

University of Utah College of Pharmacy Willard Eccles and Kuramoto fellowships (G.A.N.)

American Foundation for Pharmaceutical Education Predoctoral Fellowship (G.A.N.)

University of Utah Undergraduate Research Opportunity Program (Z.Y.)

The ALSAM Foundation (J.B.)

## Data Availability

All data generated in this study are available in this article. Densitometry-based quantitation data are available as an excel spreadsheet in **Supplemental Data File 1**. Uncropped versions of all images (including replicates) from this study are available in the **Supporting Information**.

## Supporting Information

This article contains supporting information.

