## Supporting Information for "Sirtuin 6 is a histone delactylase"

**Sirtuin 6 is a histone lysine delactylase (Supporting Information)**

#### Supporting Information

Table S1: Michaelis-Menten Kinetics of Sirt6 on Singly-Modified Nucleosomes 2

Figure S1: *In vitro* site-specific deacylation patterns of Sirt6 3

Figure S2: *In vitro* Sirt6 and Sirt7 assays 4

Figure S3: Treatment of U2OS cells with lactate and oxamate 5

Figure S4: Site-specific deacylation patterns of Sirt6 and Class I HDACs 6

Figure S5: Characterization of recombinant proteins, CRISPR knockouts, and nucleosomes 7

Figure S6: MS and HPLC analysis of MESNa peptides for histone semisynthesis 8

Figure S7: MS and HPLC analysis of semisynthetic histones 9

Sequences 10

**Uncropped Blots**

- *In Vitro* Sirt6 delactylation assay (figure 1) 11
- *In Vitro* Sirt6 Kinetics assays (figures 1 and S2) 11
- Native nucleosome site-specific deacylation (figure S1) 16
- Lactate Treatment of U2OS cells (figure 2) 17
- Treatment of U2OS cells with Lactate and Oxamate (figure S3) 20
- Treatment of U2OS cells with Rotenone (figure 3) 21
- Site-specific PTM analysis of histones from U2OS cells (figures 4 and S4) 22
- Sirt6 Overexpression (figure 5) 25
- Treatment of U2OS cells with Lactate and Panobinostat (figures 6 and S4) 26
- *In Vitro* Sirt7 delactylation assay on H3K18 nucleosomes (figure S2) 27
- Quality Control Gels (figure S5) 28

**Table S1: Michaelis-Menten Kinetics of Sirt6 on Singly Modified Mononucleosomes**

| PTM | $V_{max, app}$ (nmol*min$^{-1}$) | $k_{cat, app}$ (min$^{-1}$) | $K_{M, app}$ (nM) | $R^2$ |
|---|---|---|---|---|
| H3K9Ac | 1.7 ± 0.6 | 0.017 ± 0.006 | 90 ± 60 | 0.86 |
| H3K18Ac | 1.6 ± 1.0 | 0.016 ± 0.010 | 200 ± 100 | 0.89 |
| H3K9La | 1.8 ± 1.0 | 0.018 ± 0.010 | 150 ± 70 | 0.85 |
| H3K18La | 1.6 ± 1.0 | 0.016 ± 0.010 | 300 ± 200 | 0.73 |

#### Sirtuin 6 is a histone lysine delactylase (Supporting Information)

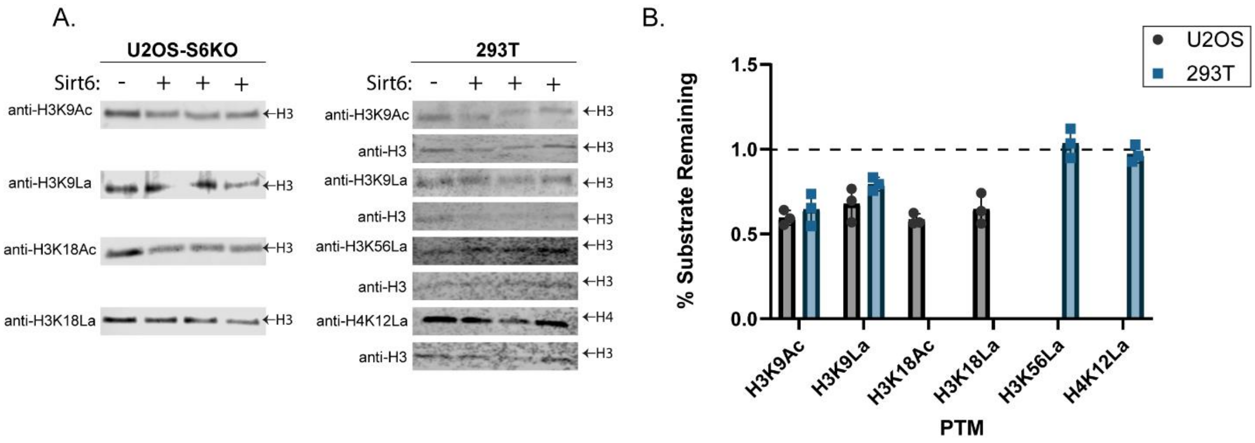


**Figure S1:** ***In vitro*** **site-specific deacylation patterns of Sirt6** (A) Nucleosomes isolated from U2OS-Sirt6 KO and HEK 293T WT cells were incubated with Sirt6 and $NAD^+$, then levels of various acyl PTMs were measured via western blot. For the 293T cells, H3 was used as a loading control. (B) Quantitation of data from (a). Data were quantified using densitometry, corrected based on a loading control (where applicable) and interpreted as a percentage of the 'untreated' control signal.

**Sirtuin 6 is a histone lysine delactylase (Supporting Information)**

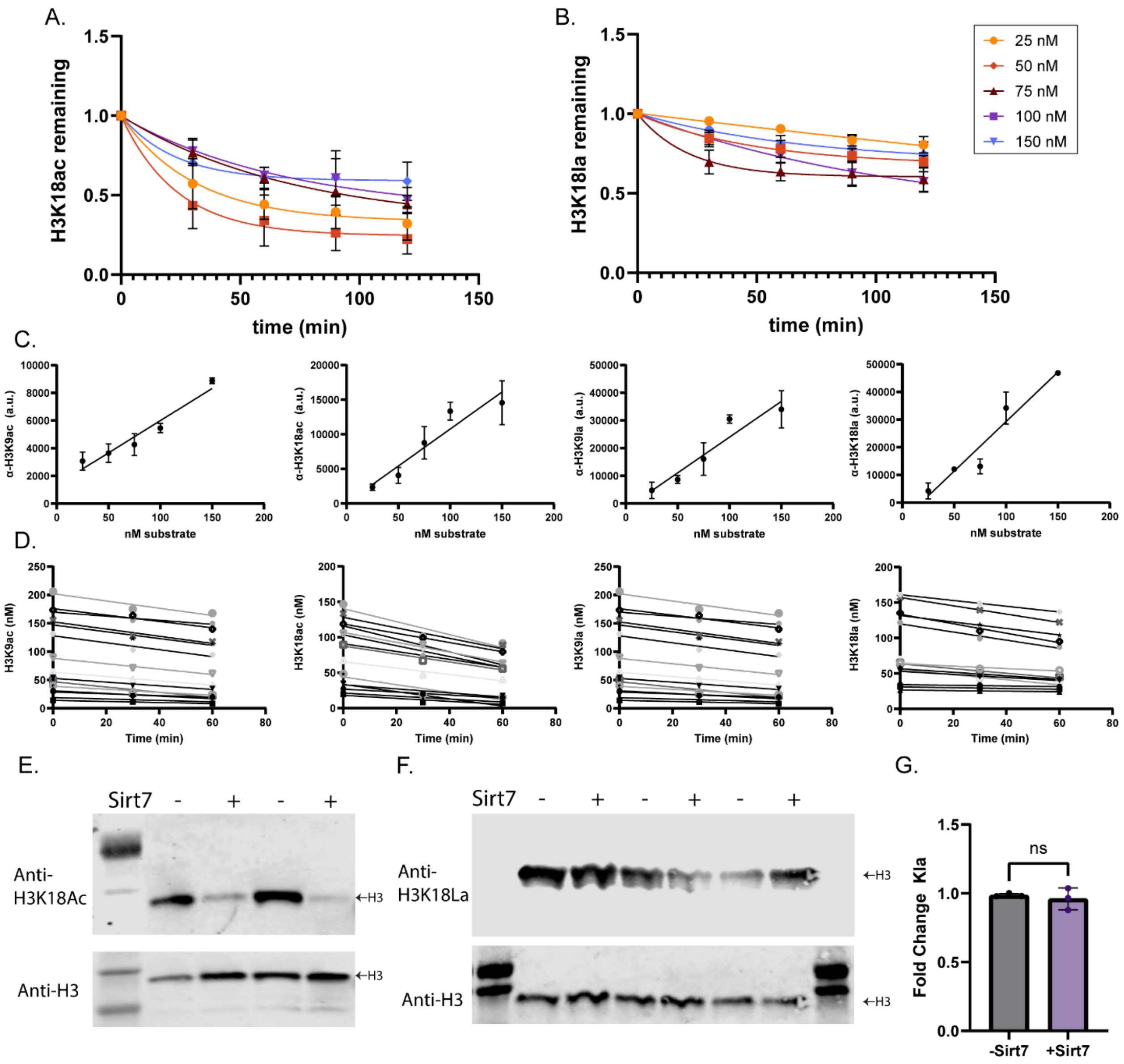


**Figure S2: *In Vitro* Sirt6 and Sirt7 assays**. (A) Sirt6 was incubated with varying concentrations of semisynthetic mononucleosome containing the H3K9ac PTM over 2 hours and the remaining substrate was measured via western blot. The data were quantified, corrected based on a loading control (histone H3 or a total protein stain), and are presented as a percentage of substrate signal remaining. The data were fit using a simple exponential decay model (y-int=1, plateau unconstrained). n=3 for each concentration and time point, error plotted as S.D. (B) as in (a), but for the H3K18ac PTM. (C) Linear range testing for the antibodies used for the kinetics assays. Data were derived from the raw fluorescence intensity and corrected based on an H3 loading control. These linear regressions were used to back-calculate the concentration of substrate remaining based on corrected intensity values from the remainder of the kinetics experiments. (D) The linear regression analysis of the first hour of data collection from the kinetics data, used to determine $V_I$ for each replicate of each condition. (E) Semisynthetic mononucleosomes (100 nM) containing the H3K18ac PTM were incubated with $NAD^+$ (1 mM) and with or without addition of Sirt7 (10 µM). Remaining H3K18ac was measured by western blot. Total protein stain (LiCor) was used as a loading control. (F) As in (d), but for the H3K18la PTM. (G) Quantitation of data from (f). Western blot data was quantified using densitometry, corrected based on the loading control, then each replicate was normalized as a fold change from the mean of the -Sirt7 condition. n=3, error plotted as S.D., p=0.60.

#### Sirtuin 6 is a histone lysine delactylase (Supporting Information)

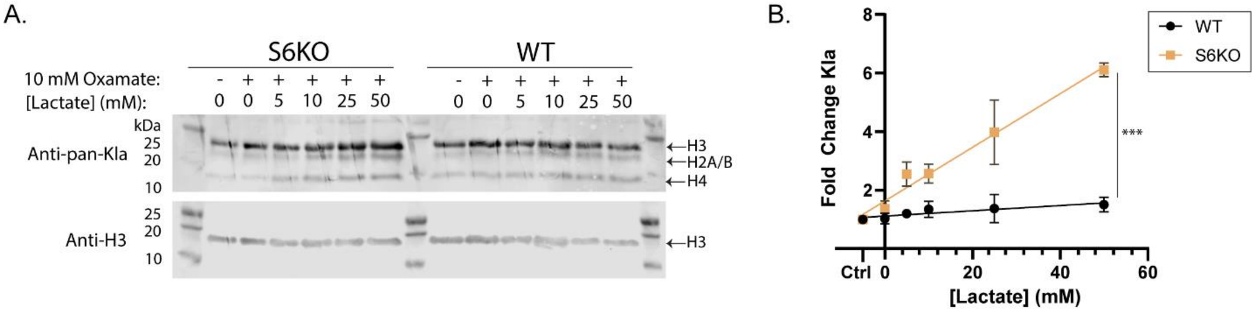


**Figure S3: Treatment of U2OS cells with lactate and oxamate**. (A) U2OS wild-type and sirt6 knockout cells were treated with oxamate and titrated with sodium lactate as indicated. Histones were acid extracted from the cells and histone Kla levels were measured using a pan-lactyllysine antibody. (B) Quantitation of data from (a). Kla signal was corrected using the H3 loading control signal, then normalized as a fold change from the untreated condition. n=3, error plotted as S.D. Slopes (starting at x=0) were compared using Welch's t-test, $p<0.0001$.

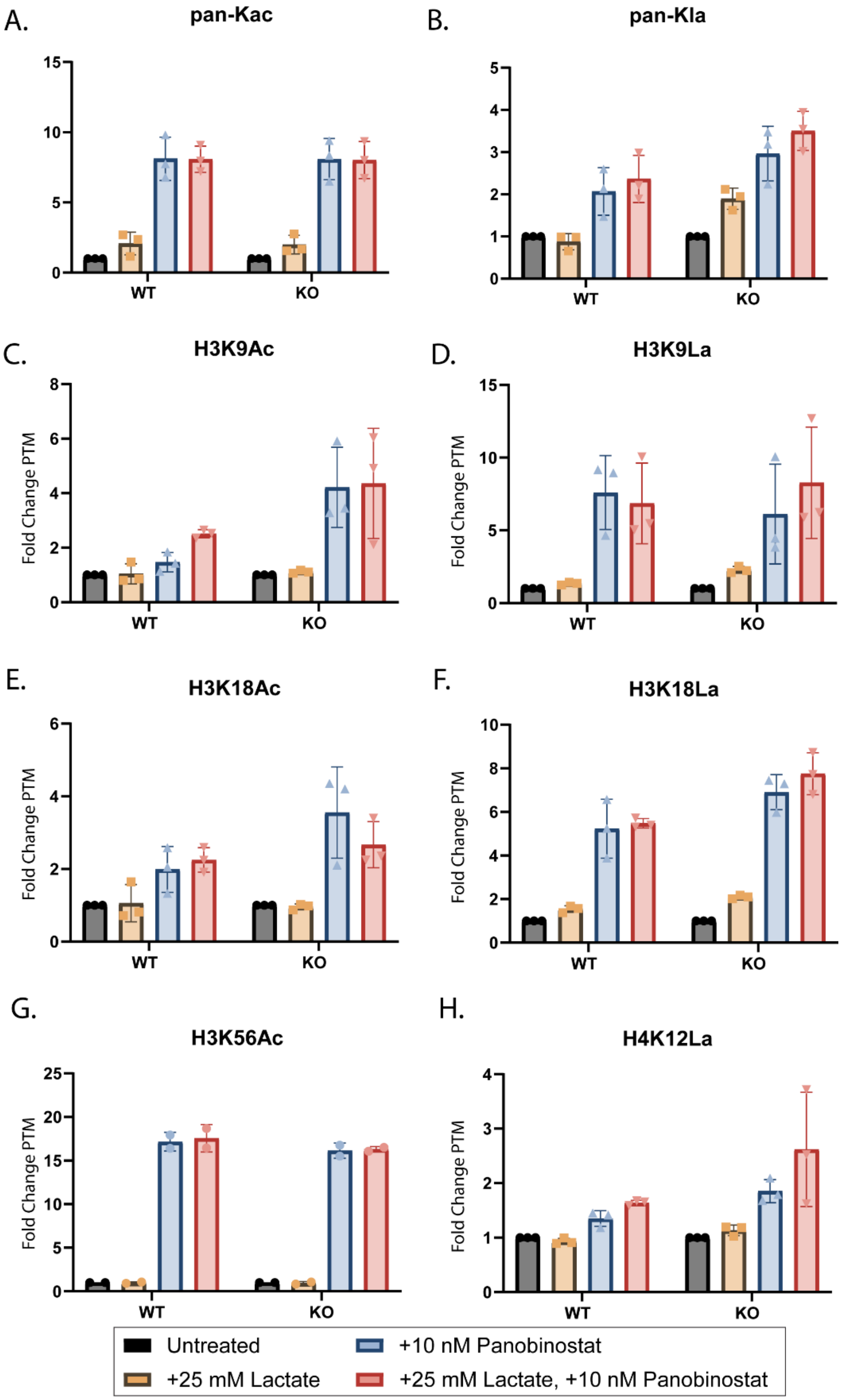


**Figure S4: Site-specific deacylation patterns of Sirt6 and Class I HDACs.** Wild-type and Sirt6-KO cells were treated with 10 nM Panobinostat, 25 mM sodium lactate, or both for 24 hours. Levels of PTMs on acid-extracted histones were measured using western blot using pan-PTM or site specific PTM antibodies (PTM Bio) as indicated. Western blot data was quantified using densitometry, corrected based on a loading control (histone H3 or total protein stain), and normalized as a fold change from the untreated condition. For (A)-(F) and (H), n=3. For (G), n=2. Error is plotted as S.D. for all panels.

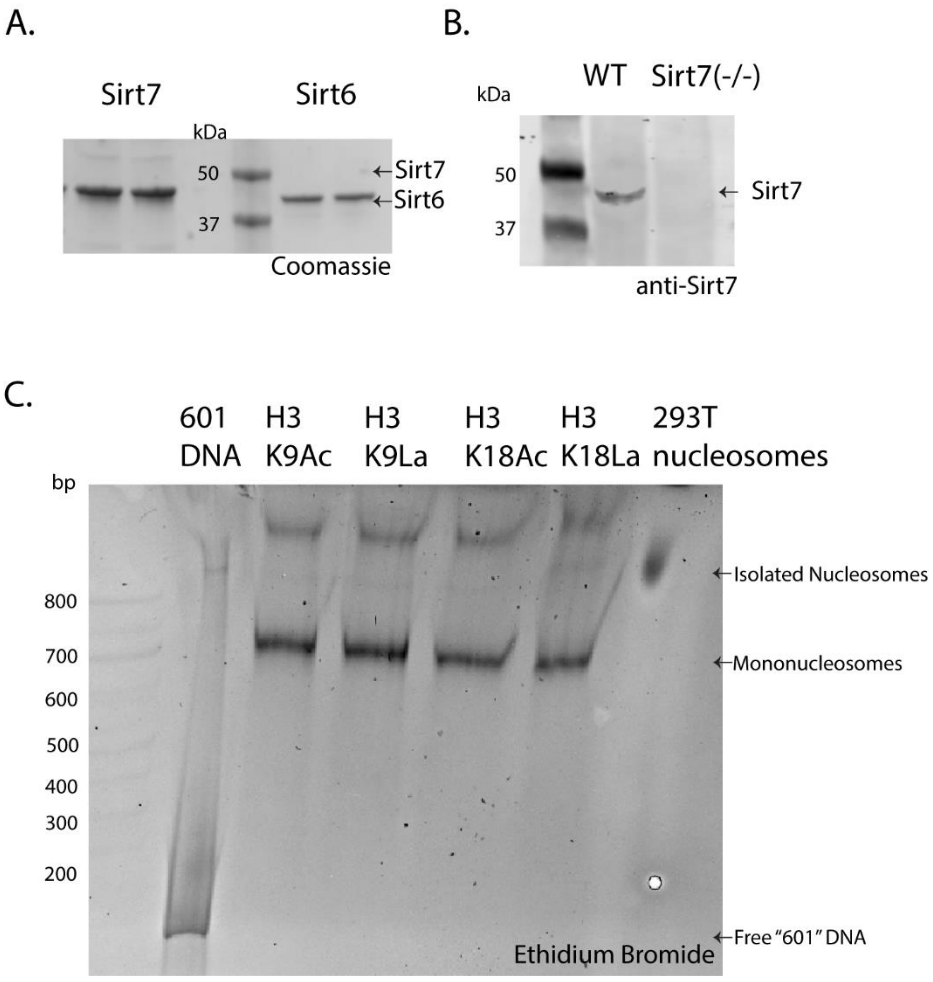


**Figure S5: Characterization of recombinant proteins, CRISPR knockouts, and nucleosomes.** (A) Coomassie stain of recombinant, full length, human Sirt6 and Sirt7 used in all *in vitro* assays. (B) Validation of Sirt7 knockout in the (-/-) Sirt7-KO cells generated using CRISPR-Cas9. Validation of Sirt6 knockout is in main-text figure 5. (C) EtBr-stained native TBE gel of all 4 semisynthetic nucleosomes and the nucleosomes isolated from 293T cells.

**Sirtuin 6 is a histone lysine delactylase (Supporting Information)**

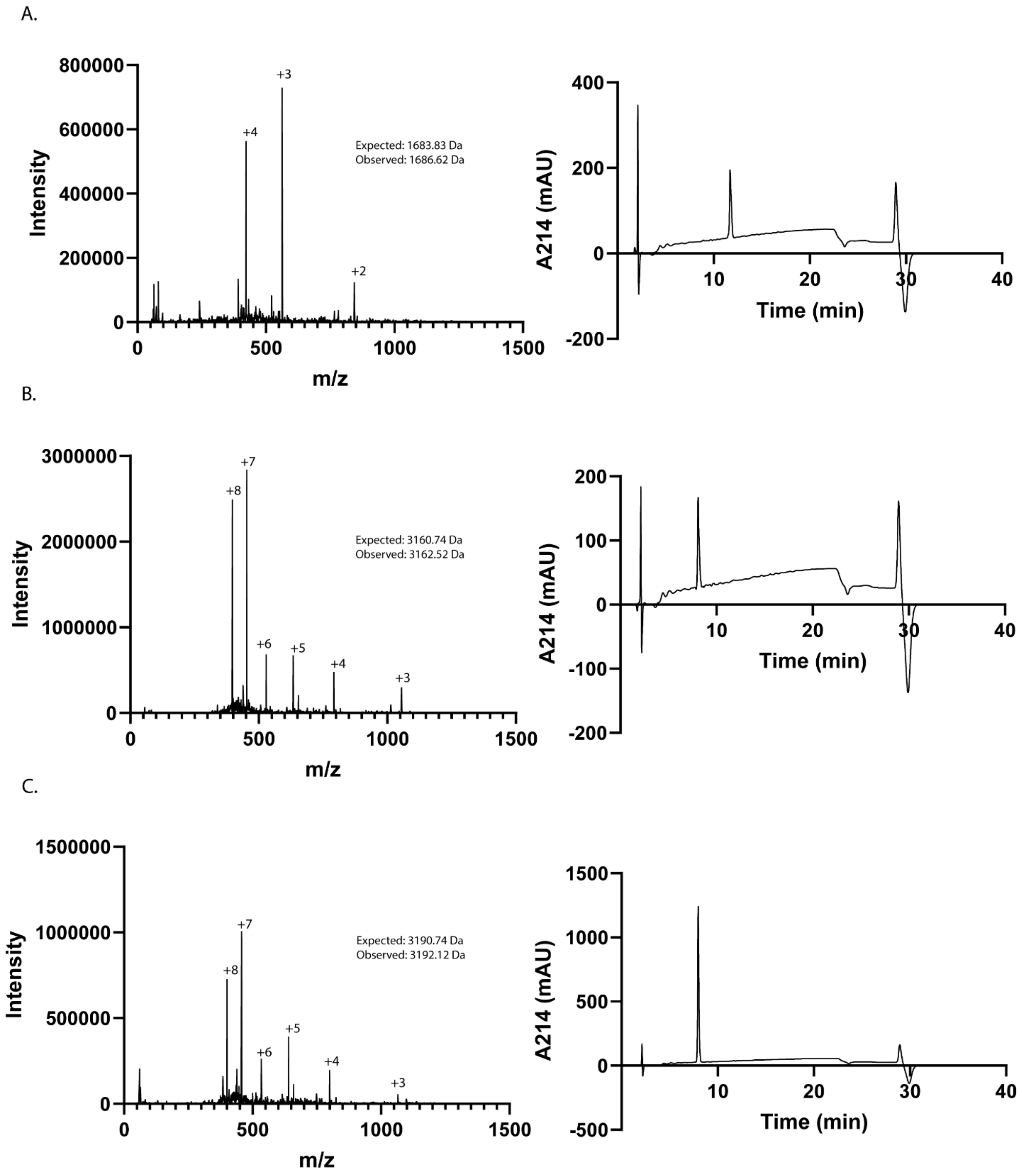


**Figure S6: MS and HPLC analysis of MESNa peptides for histone semisynthesis.** (A) LC-MS data (Waters QDa) and HPLC trace data (Waters XBridge BEH C18 analytical column) for H3K9la-MESNa peptide. (B) Data for H3K18ac-MESNa peptide. (C) Data for H3K18la-MESNa peptide. All HPLC traces collected using a 0-70% gradient of acetonitrile in water/0.1% TFA over 20 minutes.

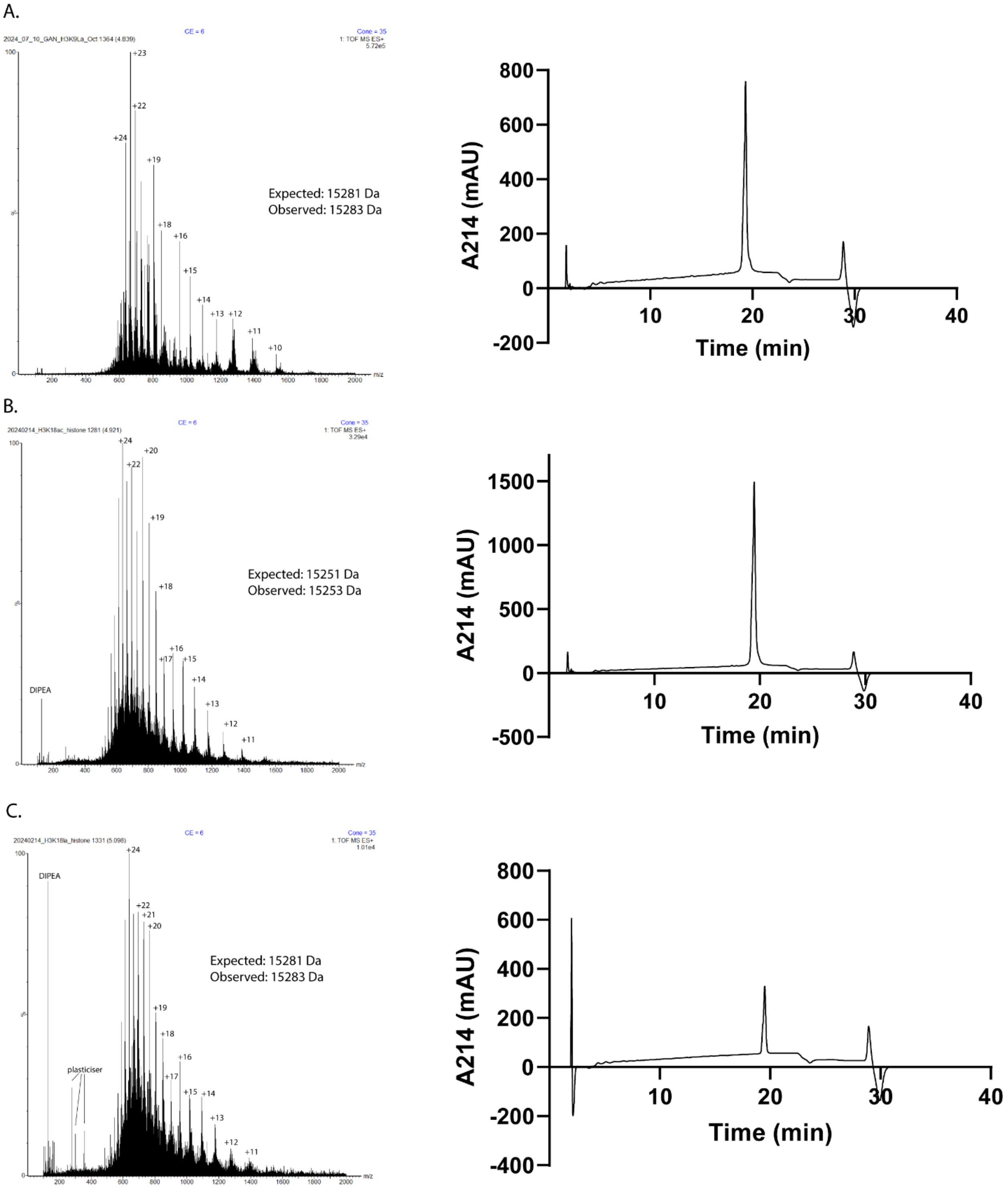


**Figure S7: MS and HPLC analysis of semisynthetic histones.** (A) High-resolution LC-MS data and HPLC trace for semisynthetic H3K9la histone. (B) Data for H3K18Ac histone. (C) Data for H3K18La histone. For all HPLC traces, separation was performed using a Waters BEH Xbridge C18 analytical column using a 0-70% gradient of acetonitrile in water/0.1% TFA over 20 min.

**Sirtuin 6 is a histone lysine delactylase (Supporting Information)**

**Sequences**

Key: **Bold = protein of interest**, underlined = SUMO tag

pET28a-LIC-6x-His-SUMO-Sirt6:

MGSSHHHHHHGSGLVPRGSASMSDSEVNQEAKPEVKPEVKPETHINLKVSDGSSEIFFKIKKTTPLRRLMEAFAKRQGKEMDSLRFLYDGIRIQADQTPEDLDMEDNDIIEAHREQIGG**SVNYAAGLSPYADKGKCGLPEIFDPPEELERKVWELARLVWQSSSVVFHTGAGISTASGIPDFRGPHGVWTMEERGLAPKFDTTFESARPTQTHMALVQLERVGLLRFLVSQNVDGLHVRSGFPRDKLAELHGNMFVEECAKCKTQYVRDTVVGTMGLKATGRLCTVAKARGLRACRGELRDTILDWEDSLPDRDLALADEASRNADLSITLGTSLQIRPSGNLPLATKRRGGRLVIVNLQPTKHDRHADLRIHGYVDEVMTRLMKHLGLEIPAWDGPRVLERALPPLPRPPTPKLEPKEESPTRINGSIPAGPKQEPCAQHNGSEPASPKRERPTSPAPHRPPKRVKAKAVPS***

pQE-80-6xHis-SUMO-Sirt7:

MRGSHHHHHHGSASDSEVNQEAKPEVKPEVKPETHINLKVSDGSSEIFFKIKKTTPLRRLMEAFAKRQGKEMDSLRFLYDGIRIQADQTPEDLDMEDNDIIEAHREQIGG**AAGGLSRSERKAAERVRRLREEQQRERLRQVCDDPEELRGKVRGLASAVRNAKYLVVYTGAGISTAASIPDYRGPNGVWTLLQKGRSVSAADLSEAEPTLTHMSITRLHEQKLVQHVVSQNCDGLHLRSGLPRTAISELHGNMYIEVCTSCVPNREYVRVFDVTERTALHRHQTGRTCHKCGTQLRDTIVHFGERGTLGQPLNWEAATEAASRADTILCLGSSLKVLKKYPRLWCMTKPPSRRPKLYIVNLQWTPKDDWAALKLHGKCDDVMRLLMAELGLEIPAYSRWQDPIFSLATPLRAGEEGSHSRKSLCRSREEAPPGDRGAPLSSAPILGGWFGRGCTKRTKRKKVT**EFGGDYKDDDDK*

pLJM1-Sirt6-3xFLAG

M**SVNYAAGLSPYADKGKCGLPEIFDPPEELERKVWELARLVWQSSSVVFHTGAGISTASGIPDFRGPHGVWTMEERGLAPKFDTTFESARPTQTHMALVQLERVGLLRFLVSQNVDGLHVRSGFPRDKLAELHGNMFVEECAKCKTQYVRDTVVGTMGLKATGRLCTVAKARGLRACRGELRDTILDWEDSLPDRDLALADEASRNADLSITLGTSLQIRPSGNLPLATKRRGGRLVIVNLQPTKHDRHADLRIHGYVDEVMTRLMKHLGLEIPAWDGPRVLERALPPLPRPPTPKLEPKEESPTRINGSIPAGPKQEPCAQHNGSEPASPKRERPTSPAPHRPPKRVKAKAVPS**EFGGDYKDDDDKGGSDYKDDDDKGGSDYKDDDDK*

ds601 DNA:

Key: 601 sequence (147 bp) in bold

CTACTGGTACGGCAG**ACAGGATGTATATATCTGACACGTGCCTGGAGACTAGGGAGTAATCCCCTTGGCGGTTAAAACGCGGGGGACAGCGCGTACGTGCGTTTAAGCGGTGCTAGAGCTGTCTACGACCAATTGAGCGGCCTCGGCACCGGGATTCTCCAG**TATTCGAGGCCGTTC

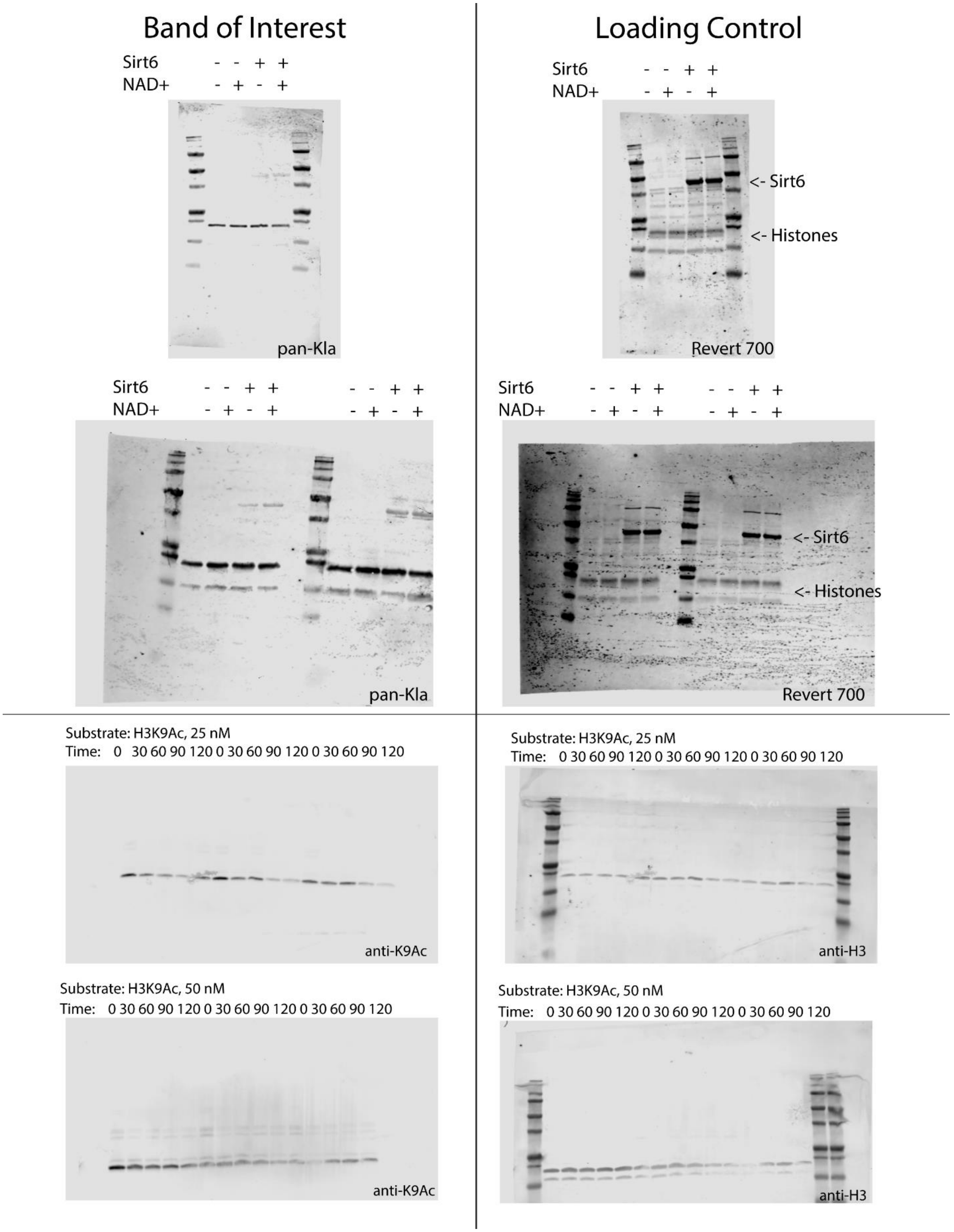
Band of Interest
Loading Control
Sirt6 - - + +
NAD+ - + - +
pan-Kla
Sirt6 - - + +
NAD+ - + - +
<- Sirt6
<- Histones
Revert 700
Sirt6 - - + + - - + +
NAD+ - + - + - + - +
pan-Kla
Sirt6 - - + + - - + +
NAD+ - + - + - + - +
<- Sirt6
<- Histones
Revert 700
Substrate: H3K9Ac, 25 nM
Time: 0 30 60 90 120 0 30 60 90 120 0 30 60 90 120
anti-K9Ac
Substrate: H3K9Ac, 25 nM
Time: 0 30 60 90 120 0 30 60 90 120 0 30 60 90 120
anti-H3
Substrate: H3K9Ac, 50 nM
Time: 0 30 60 90 120 0 30 60 90 120 0 30 60 90 120
anti-K9Ac
Substrate: H3K9Ac, 50 nM
Time: 0 30 60 90 120 0 30 60 90 120 0 30 60 90 120
anti-H3

**Sirtuin 6 is a histone lysine delactylase (Supporting Information)**

### Band of Interest

Substrate: H3K9Ac, 75 nM
Time: 0 30 60 90 120 0 30 60 90 120 0 30 60 90 120

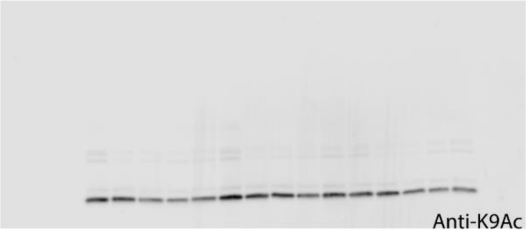


Substrate: H3K9Ac, 100 nM
Time: 0 30 60 90 120 0 30 60 90 120 0 30 60 90 120

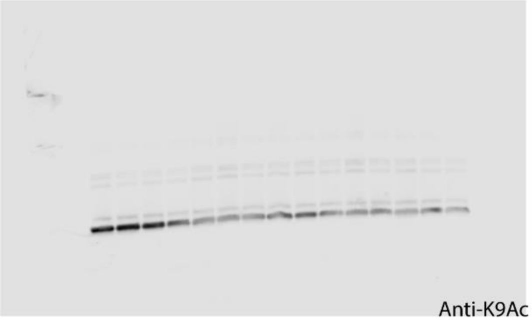


Substrate: H3K9Ac, 150 nM
Time: 0 30 60 90 120 0 30 60 90 120 0 30 60 90 120

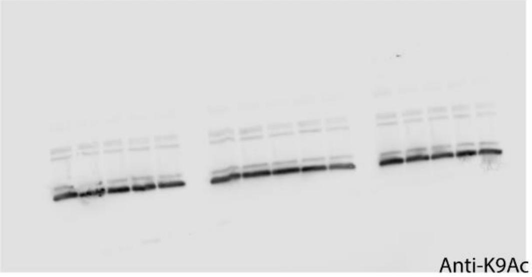


Substrate: H3K9La, 25 nM
Time: 0 30 60 90 120 0 30 60 90 120 0 30 60 90 120

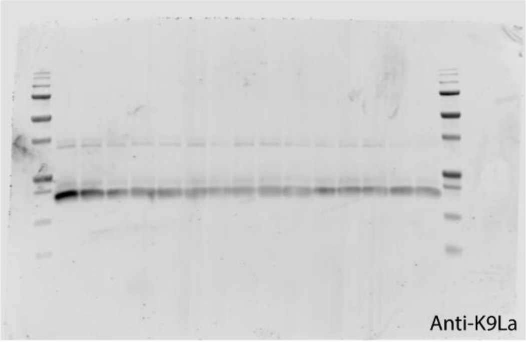


### Loading Control

Substrate: H3K9Ac, 75 nM
Time: 0 30 60 90 120 0 30 60 90 120 0 30 60 90 120

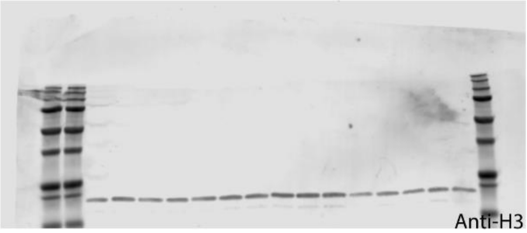


Substrate: H3K9Ac, 100 nM
Time: 0 30 60 90 120 0 30 60 90 120 0 30 60 90 120

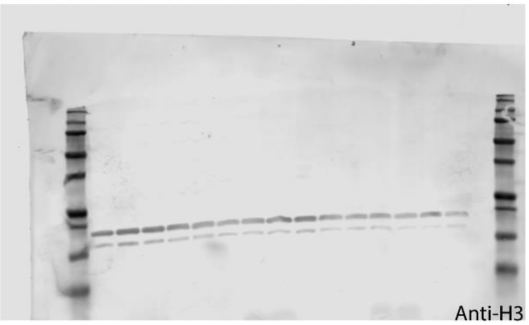


Substrate: H3K9Ac, 150 nM
Time: 0 30 60 90 120 0 30 60 90 120 0 30 60 90 120

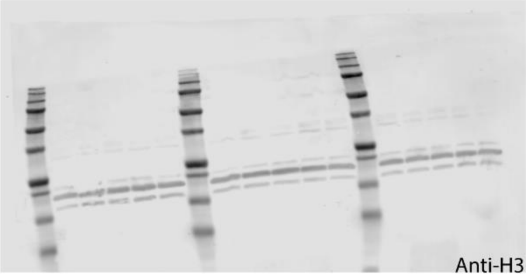


Substrate: H3K9La, 25 nM
Time: 0 30 60 90 120 0 30 60 90 120 0 30 60 90 120

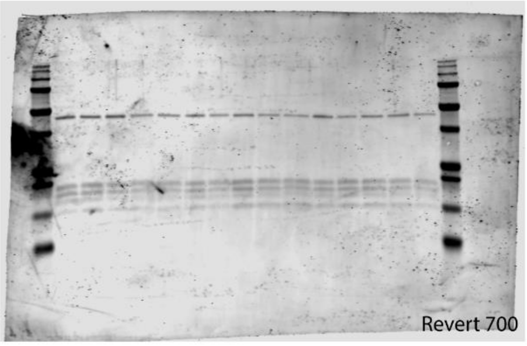

**Sirtuin 6 is a histone lysine delactylase (Supporting Information)**

### Band of Interest

Substrate: H3K9La, 50 nM
Time: 0 30 60 90 120 0 30 60 90 120 0 30 60 90 120

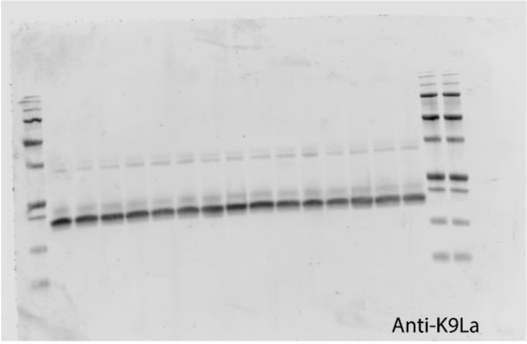


Substrate: H3K9La, 75 nM
Time: 0 30 60 90 120 0 30 60 90 120 0 30 60 90 120

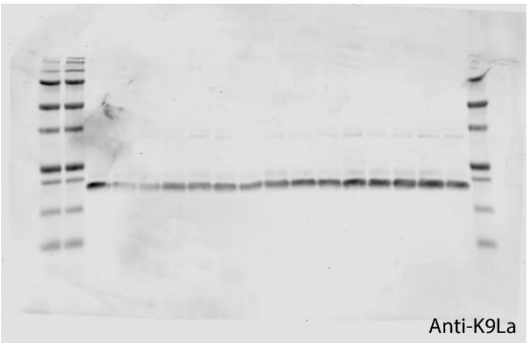


Substrate: H3K9La, 100 nM
Time: 0 30 60 90 120 0 30 60 90 120 0 30 60 90 120

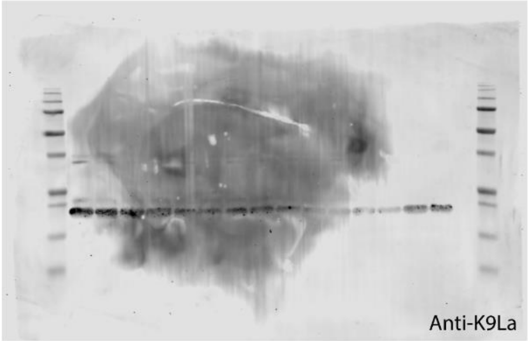


Substrate: H3K9La, 150 nM
Time: 0 30 60 90 120 0 30 60 90 120 0 30 60 90 120

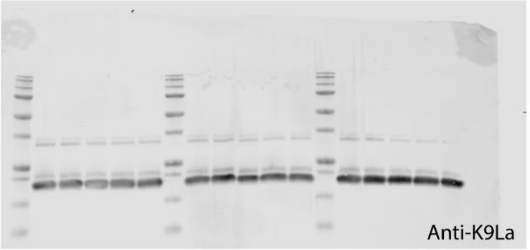


### Loading Control

Substrate: H3K9La, 50 nM
Time: 0 30 60 90 120 0 30 60 90 120 0 30 60 90 120

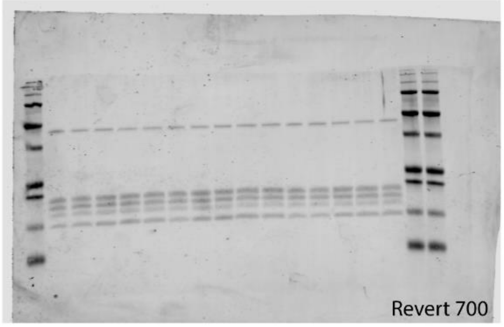


Substrate: H3K9La, 75 nM
Time: 0 30 60 90 120 0 30 60 90 120 0 30 60 90 120

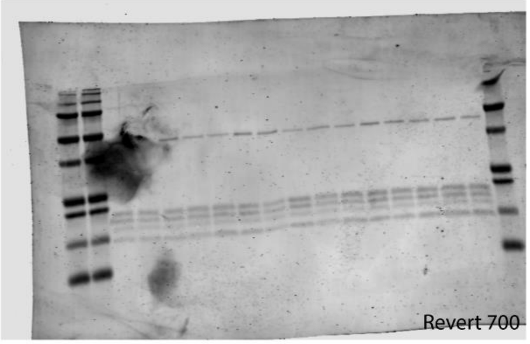


Substrate: H3K9La, 100 nM
Time: 0 30 60 90 120 0 30 60 90 120 0 30 60 90 120

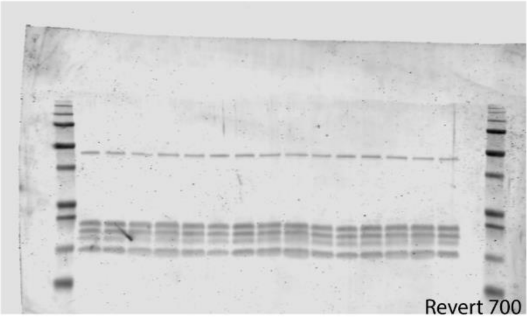


Substrate: H3K9La, 150 nM
Time: 0 30 60 90 120 0 30 60 90 120 0 30 60 90 120

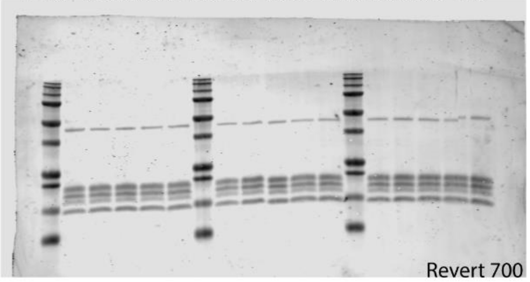

**Sirtuin 6 is a histone lysine delactylase (Supporting Information)**

#### Band of Interest

Substrate: H3K18Ac, 25nM
Time: 0 30 60 90 120 0 30 60 90 120 0 30 60 90 120

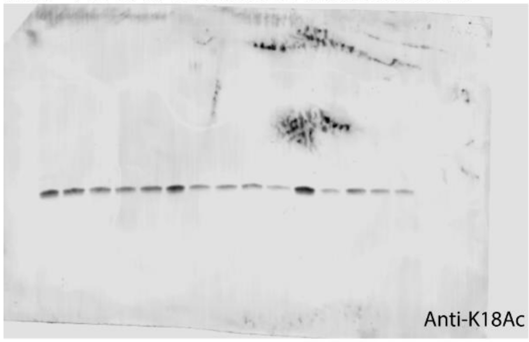


Substrate: H3K18Ac, 50 nM
Time: 0 30 60 90 120 0 30 60 90 120 0 30 60 90 120

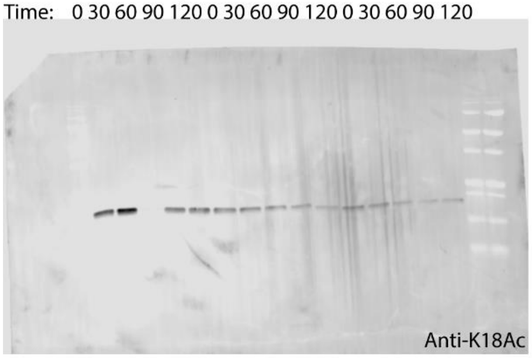


Substrate: H3K18Ac, 75 nM
Time: 0 30 60 90 120 0 30 60 90 120 0 30 60 90 120

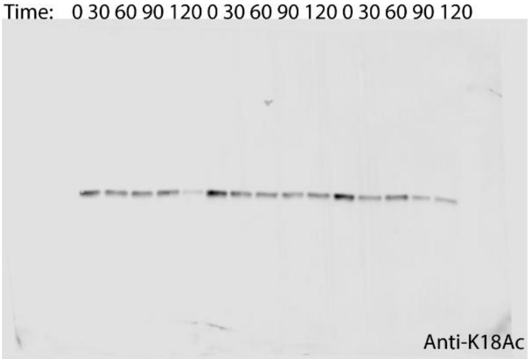


Substrate: H3K18Ac, 100 nM
Time: 0 30 60 90 120 0 30 60 90 120 0 30 60 90 120

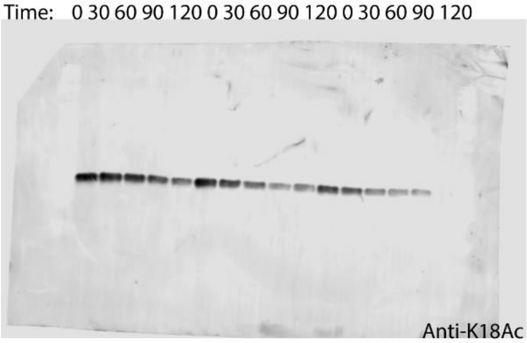


#### Loading Control

Substrate: H3K18Ac, 25nM
Time: 0 30 60 90 120 0 30 60 90 120 0 30 60 90 120

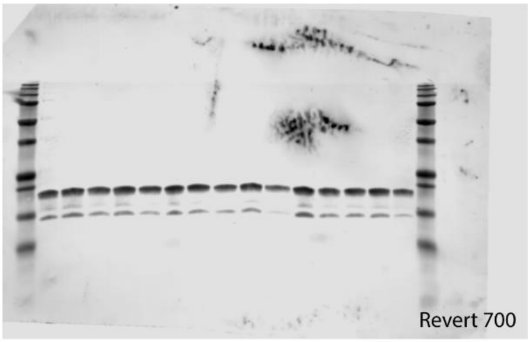


Substrate: H3K18Ac, 50 nM
Time: 0 30 60 90 120 0 30 60 90 120 0 30 60 90 120

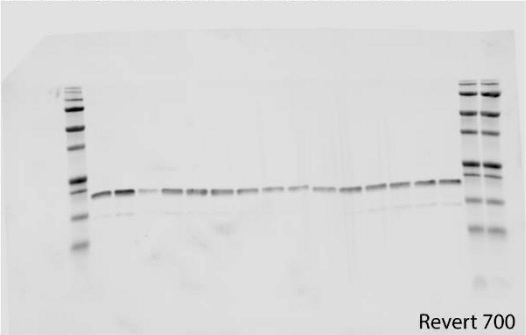


Substrate: H3K18Ac, 75nM
Time: 0 30 60 90 120 0 30 60 90 120 0 30 60 90 120

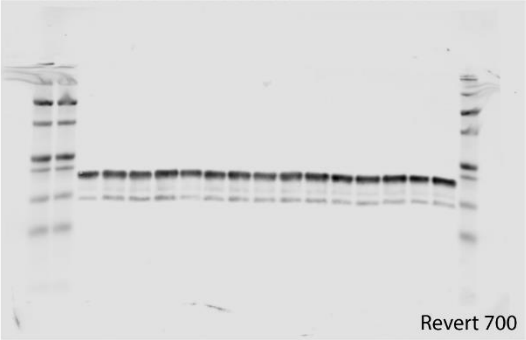


Substrate: H3K18Ac, 100nM
Time: 0 30 60 90 120 0 30 60 90 120 0 30 60 90 120

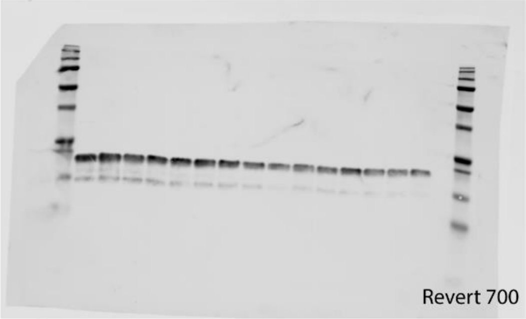

#### Band of Interest

Substrate: H3K18Ac, 150 nM
Time: 0 30 60 90 120 0 30 60 90 120 0 30 60 90 120

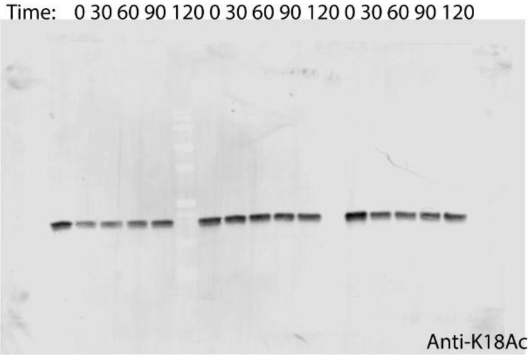


Substrate: H3K18La, 25 nM
Time: 0 30 60 90 120 0 30 60 90 120 0 30 60 90 120

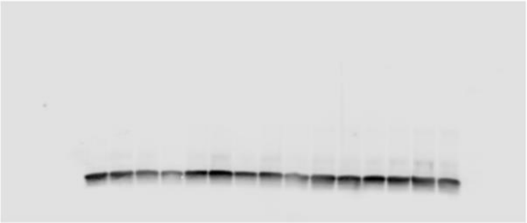

Anti-K18La

Substrate: H3K18La, 50 nM
Time: 0 30 60 90 120 0 30 60 90 120 0 30 60 90 120

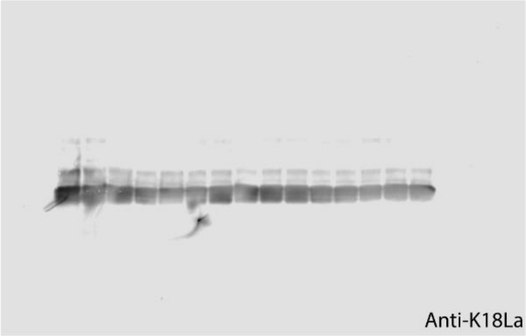


Substrate: H3K18La, 75 nM
Time: 0 30 60 90 120 0 30 60 90 120 0 30 60 90 120

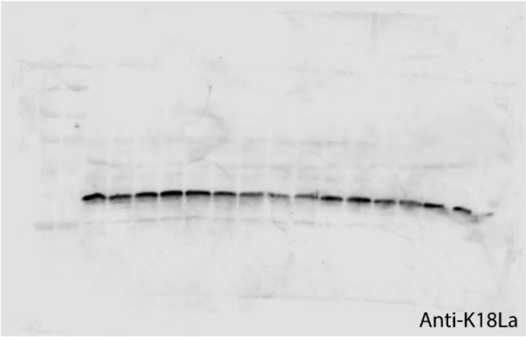


#### Loading Control

Substrate: H3K18Ac, 150 nM
Time: 0 30 60 90 120 0 30 60 90 120 0 30 60 90 120

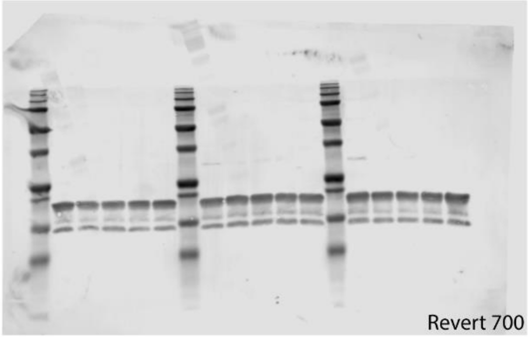


Substrate: H3K18La, 25 nM
Time: 0 30 60 90 120 0 30 60 90 120 0 30 60 90 120

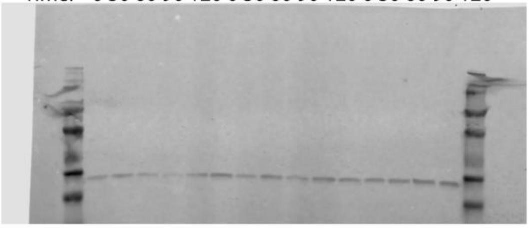

anti-H3

Substrate: H3K18La, 50 nM
Time: 0 30 60 90 120 0 30 60 90 120 0 30 60 90 120

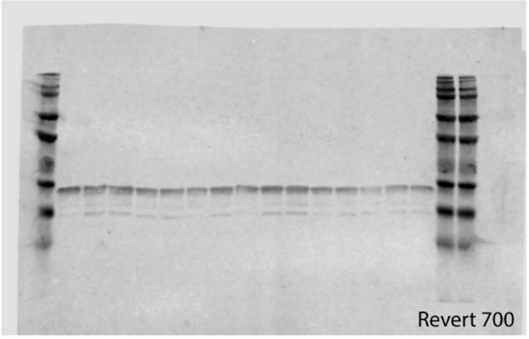


Substrate: H3K18La, 75 nM
Time: 0 30 60 90 120 0 30 60 90 120 0 30 60 90 120

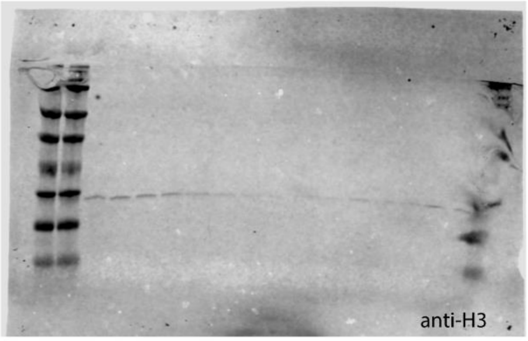

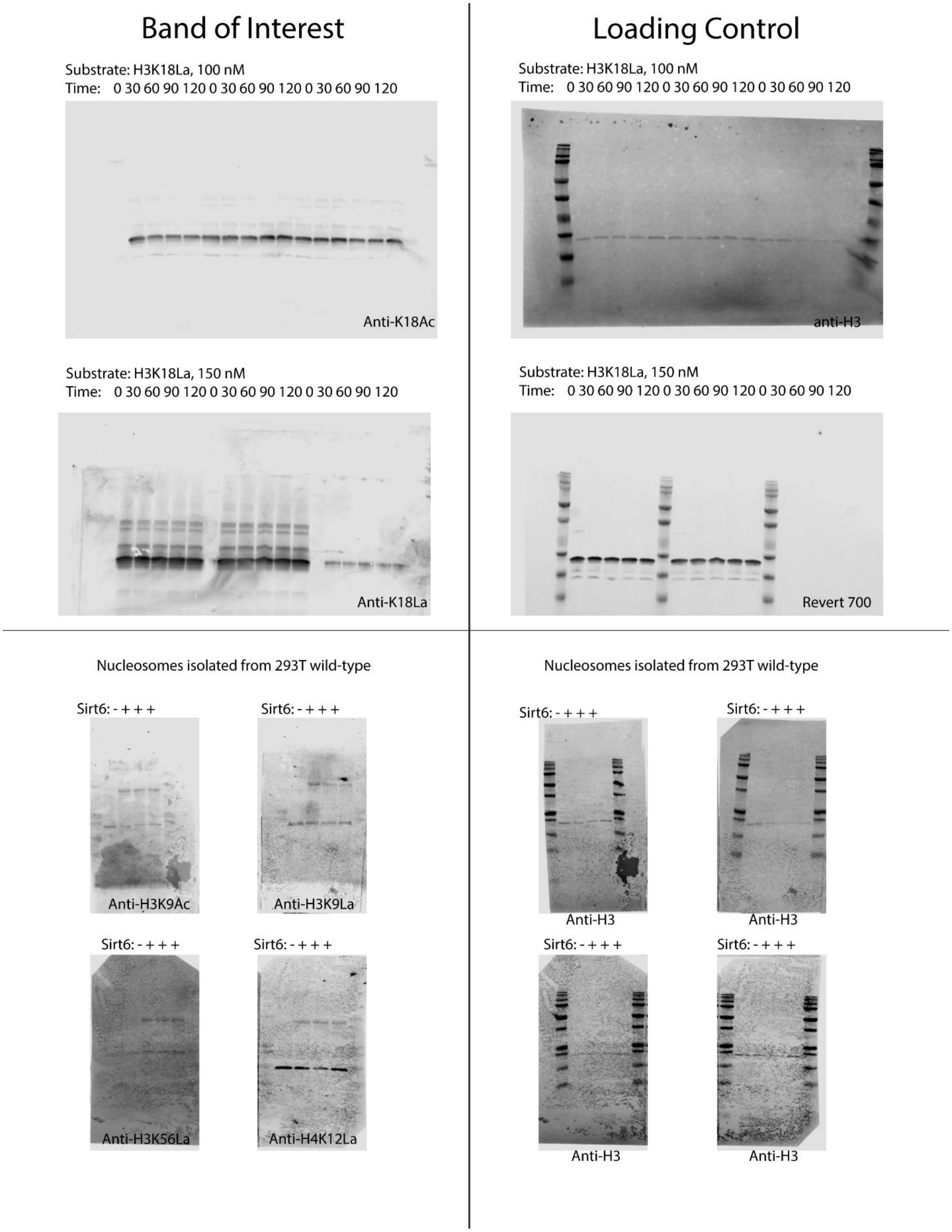
Band of Interest
Loading Control
Substrate: H3K18La, 100 nM
Time: 0 30 60 90 120 0 30 60 90 120 0 30 60 90 120
Anti-K18Ac
Substrate: H3K18La, 100 nM
Time: 0 30 60 90 120 0 30 60 90 120 0 30 60 90 120
anti-H3
Substrate: H3K18La, 150 nM
Time: 0 30 60 90 120 0 30 60 90 120 0 30 60 90 120
Anti-K18La
Substrate: H3K18La, 150 nM
Time: 0 30 60 90 120 0 30 60 90 120 0 30 60 90 120
Revert 700
Nucleosomes isolated from 293T wild-type
Sirt6: - + + +
Anti-H3K9Ac
Sirt6: - + + +
Anti-H3K9La
Sirt6: - + + +
Anti-H3K56La
Sirt6: - + + +
Anti-H4K12La
Nucleosomes isolated from 293T wild-type
Sirt6: - + + +
Anti-H3
Sirt6: - + + +
Anti-H3
Sirt6: - + + +
Anti-H3
Sirt6: - + + +
Anti-H3

**Sirtuin 6 is a histone lysine delactylase (Supporting Information)**

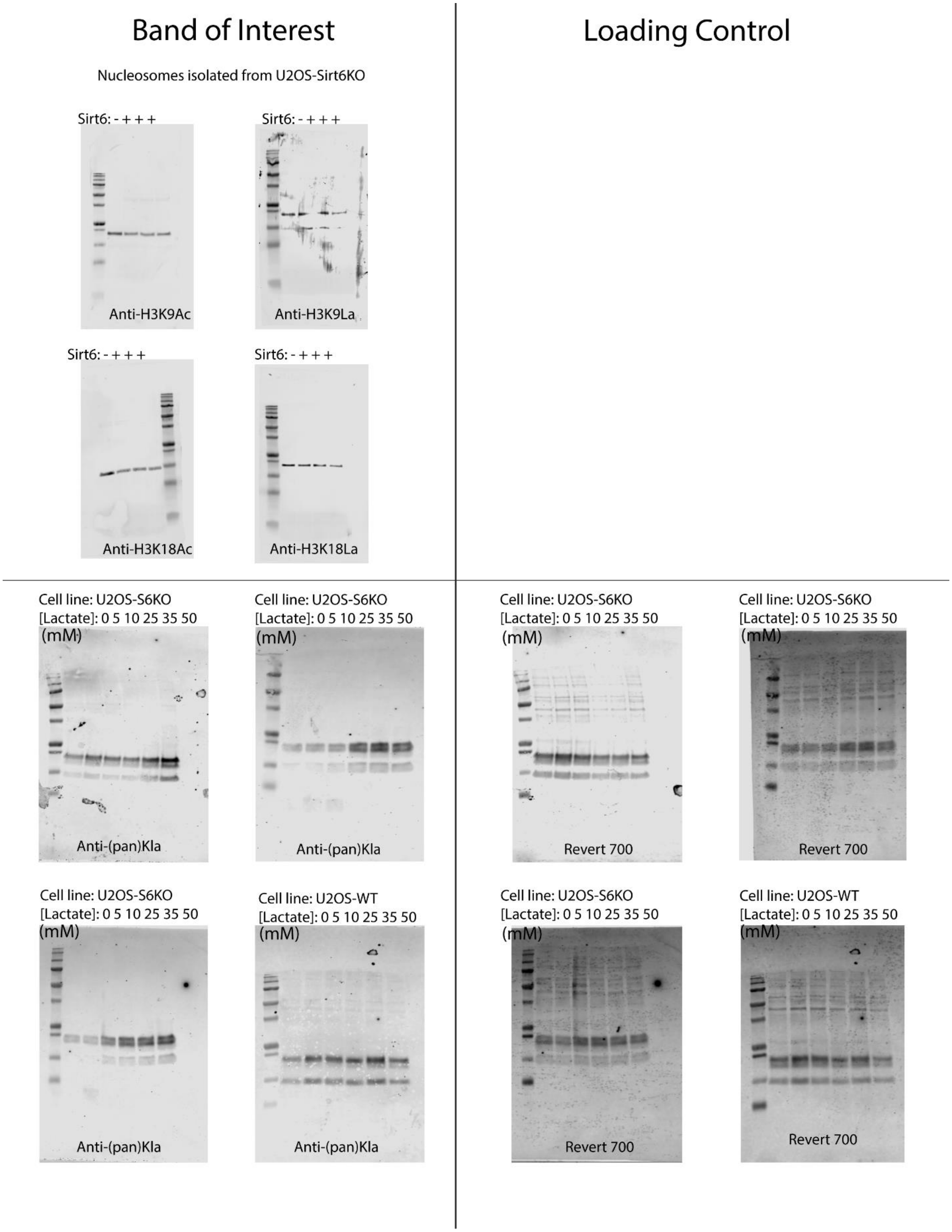

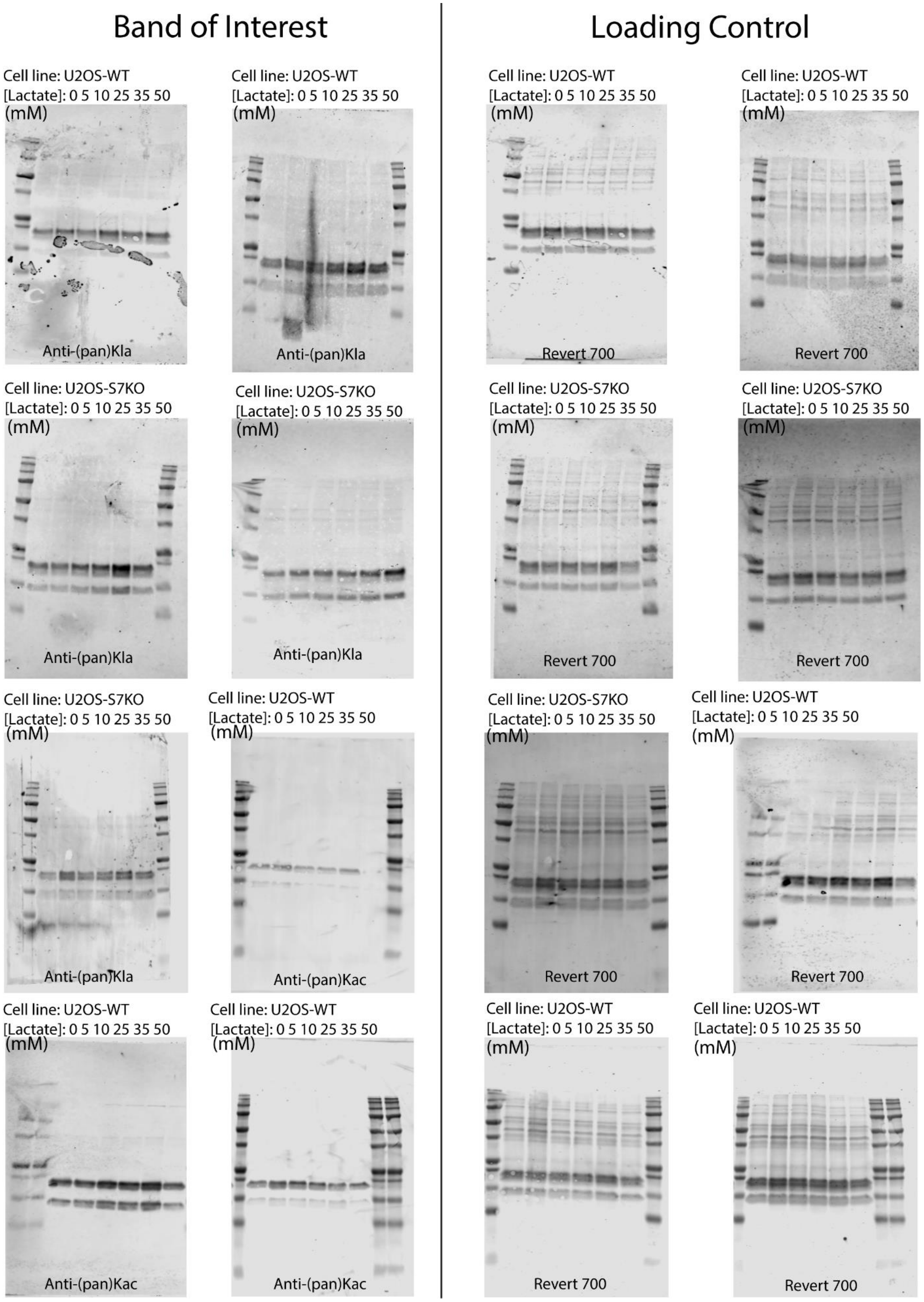
Band of Interest
Loading Control
Cell line: U2OS-WT
[Lactate]: 0 5 10 25 35 50
(mM)
Anti-(pan)Kla
Cell line: U2OS-WT
[Lactate]: 0 5 10 25 35 50
(mM)
Anti-(pan)Kla
Cell line: U2OS-WT
[Lactate]: 0 5 10 25 35 50
(mM)
Revert 700
Cell line: U2OS-WT
[Lactate]: 0 5 10 25 35 50
(mM)
Revert 700
Cell line: U2OS-S7KO
[Lactate]: 0 5 10 25 35 50
(mM)
Anti-(pan)Kla
Cell line: U2OS-S7KO
[Lactate]: 0 5 10 25 35 50
(mM)
Anti-(pan)Kla
Cell line: U2OS-S7KO
[Lactate]: 0 5 10 25 35 50
(mM)
Revert 700
Cell line: U2OS-S7KO
[Lactate]: 0 5 10 25 35 50
(mM)
Revert 700
Cell line: U2OS-S7KO
[Lactate]: 0 5 10 25 35 50
(mM)
Anti-(pan)Kla
Cell line: U2OS-WT
[Lactate]: 0 5 10 25 35 50
(mM)
Anti-(pan)Kac
Cell line: U2OS-S7KO
[Lactate]: 0 5 10 25 35 50
(mM)
Revert 700
Cell line: U2OS-WT
[Lactate]: 0 5 10 25 35 50
(mM)
Revert 700
Cell line: U2OS-WT
[Lactate]: 0 5 10 25 35 50
(mM)
Anti-(pan)Kac
Cell line: U2OS-WT
[Lactate]: 0 5 10 25 35 50
(mM)
Anti-(pan)Kac
Cell line: U2OS-WT
[Lactate]: 0 5 10 25 35 50
(mM)
Revert 700
Cell line: U2OS-WT
[Lactate]: 0 5 10 25 35 50
(mM)
Revert 700

**Sirtuin 6 is a histone lysine delactylase (Supporting Information)**

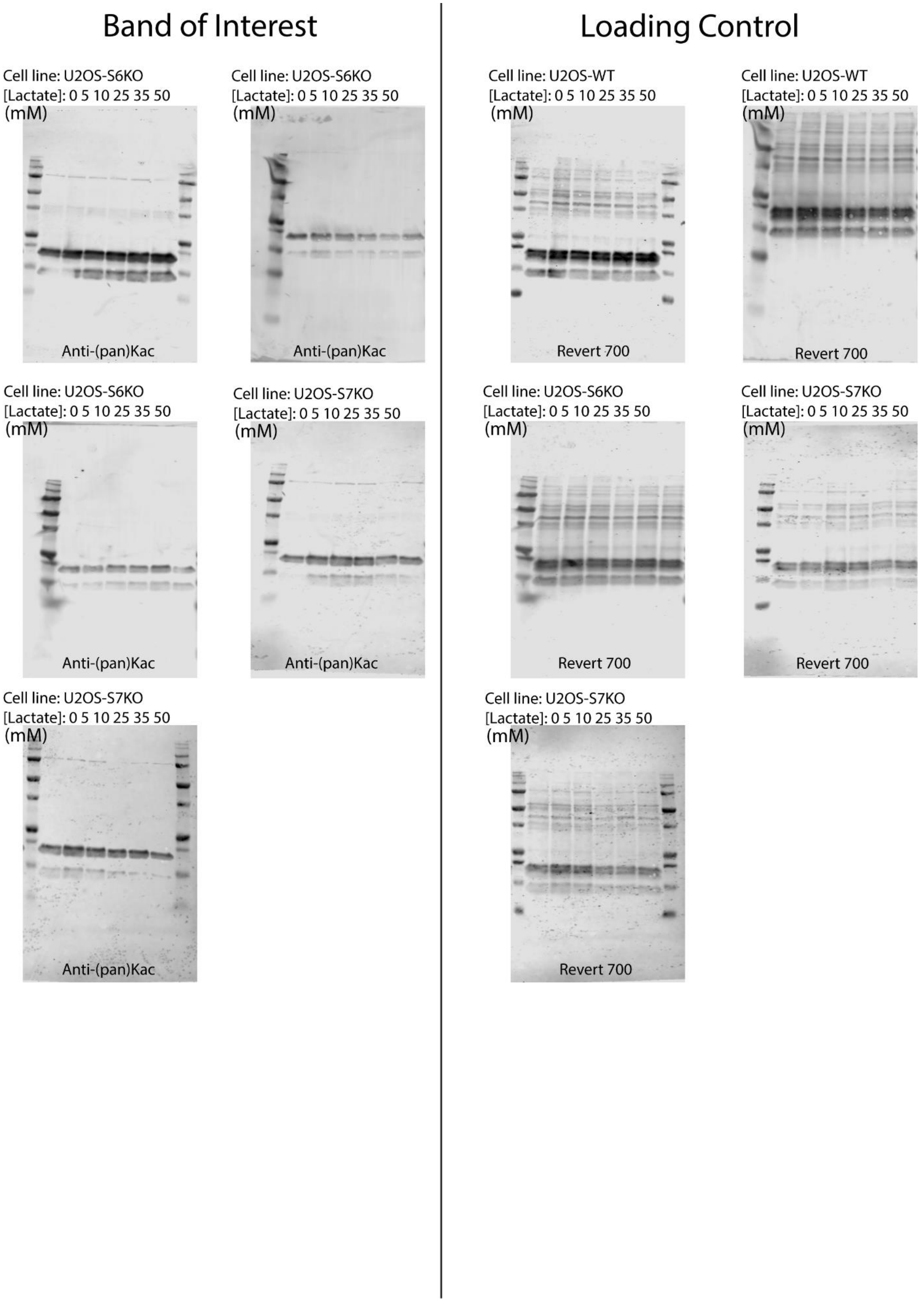

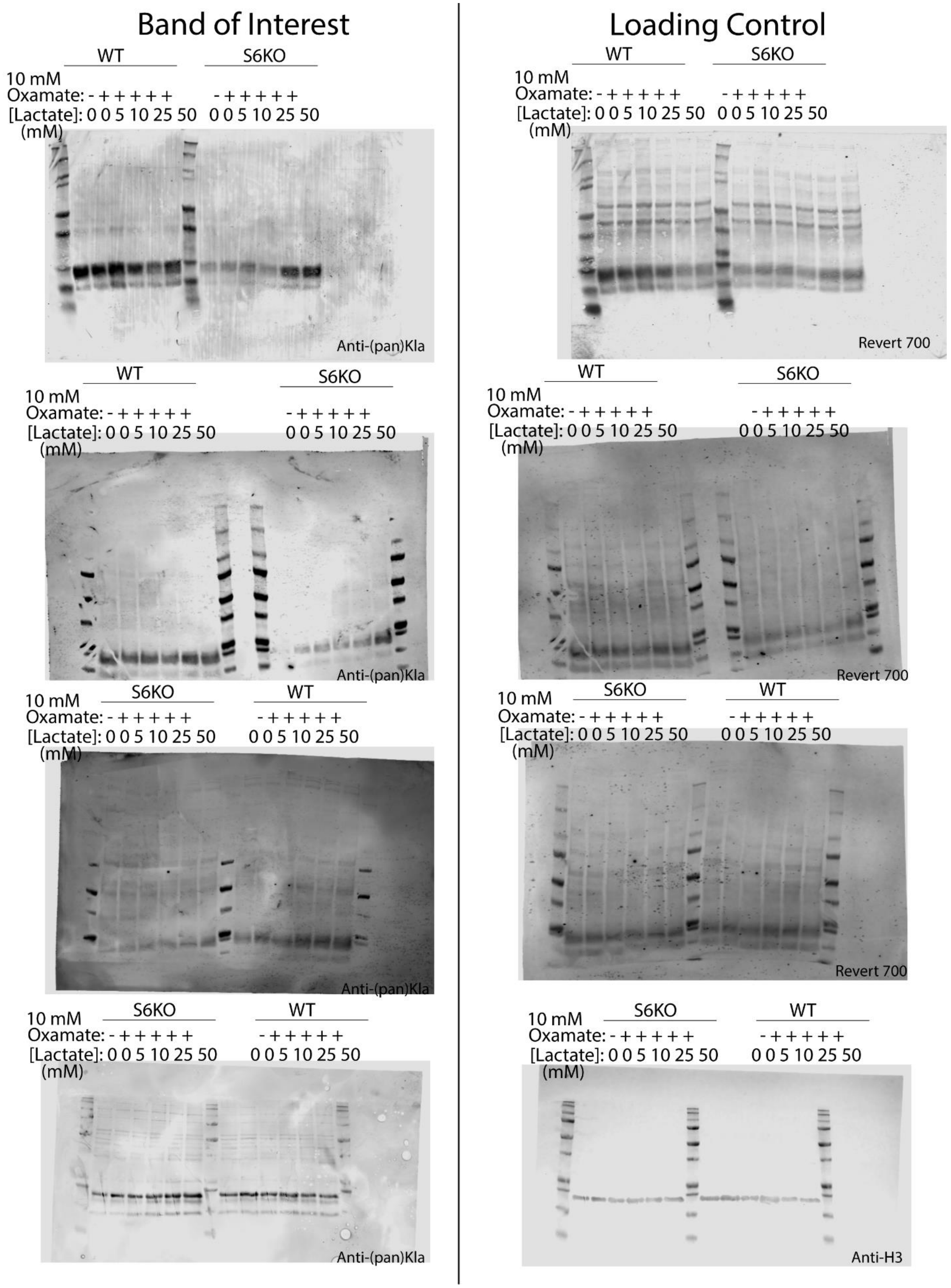
Band of Interest
Loading Control
WT
S6KO
10 mM
Oxamate: - + + + + +
[Lactate]: 0 0 5 10 25 50
(mM)
Anti-(pan)Kla
Revert 700
Anti-H3

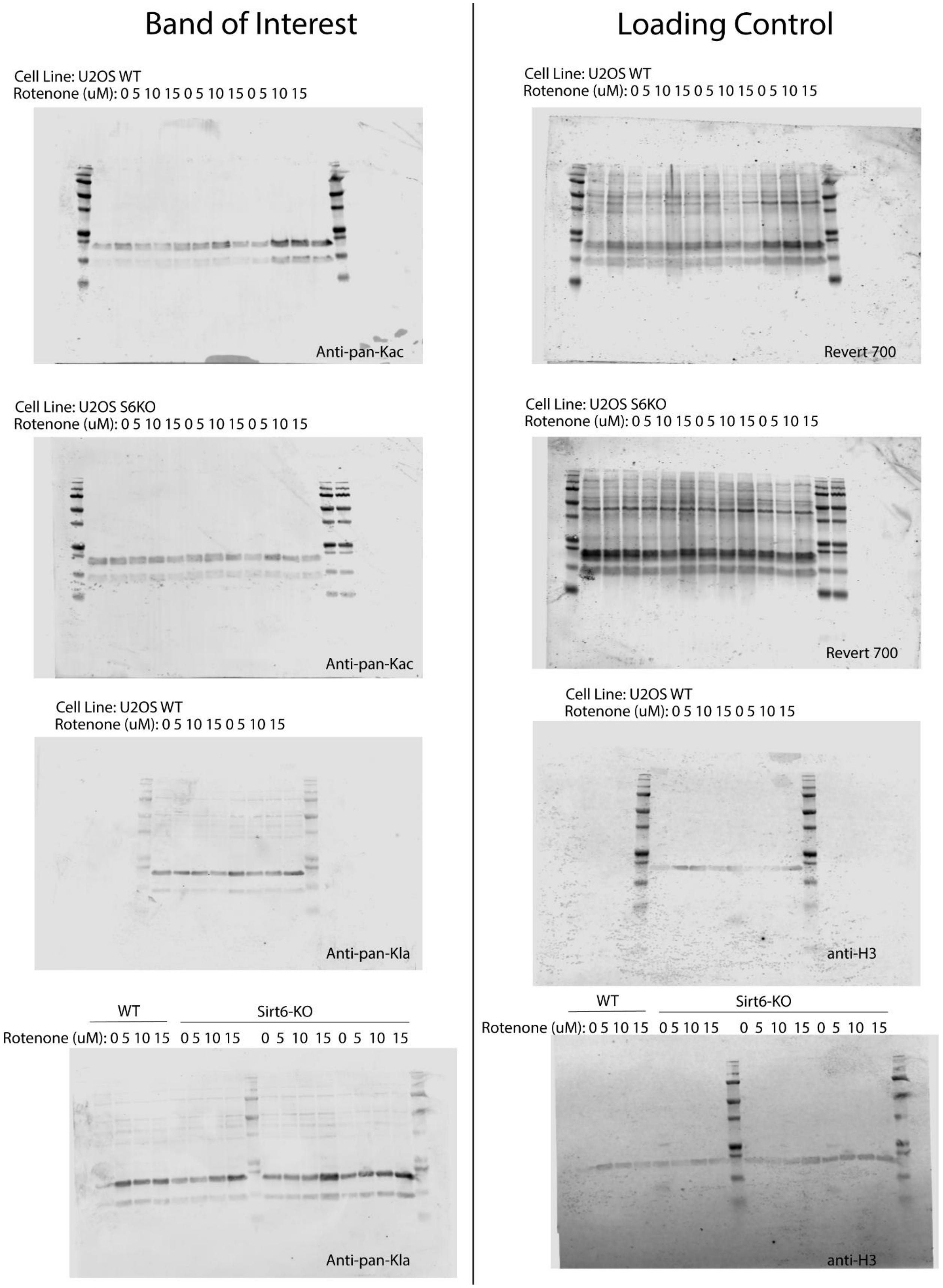
Band of Interest
Loading Control
Cell Line: U2OS WT
Rotenone (uM): 0 5 10 15 0 5 10 15 0 5 10 15
Anti-pan-Kac
Cell Line: U2OS WT
Rotenone (uM): 0 5 10 15 0 5 10 15 0 5 10 15
Revert 700
Cell Line: U2OS S6KO
Rotenone (uM): 0 5 10 15 0 5 10 15 0 5 10 15
Anti-pan-Kac
Cell Line: U2OS S6KO
Rotenone (uM): 0 5 10 15 0 5 10 15 0 5 10 15
Revert 700
Cell Line: U2OS WT
Rotenone (uM): 0 5 10 15 0 5 10 15
Anti-pan-Kla
Cell Line: U2OS WT
Rotenone (uM): 0 5 10 15 0 5 10 15
anti-H3
WT
Sirt6-KO
Rotenone (uM): 0 5 10 15 0 5 10 15 0 5 10 15 0 5 10 15
Anti-pan-Kla
WT
Sirt6-KO
Rotenone (uM): 0 5 10 15 0 5 10 15 0 5 10 15 0 5 10 15
anti-H3

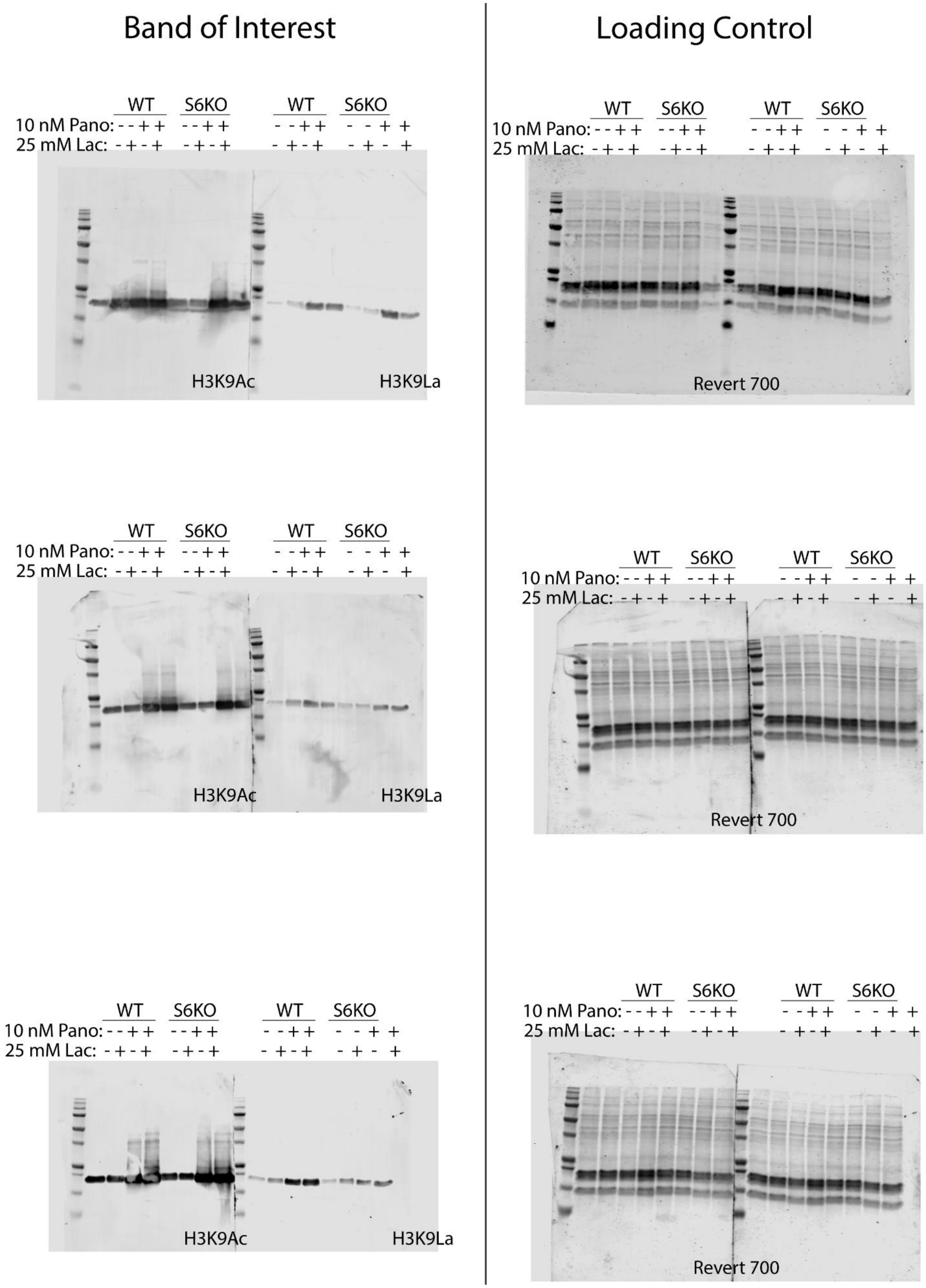
Band of Interest
Loading Control
WT S6KO WT S6KO
10 nM Pano: - - + + - - + + - - + + - - + +
25 mM Lac: - + - + - + - + - + - + - + - +
H3K9Ac
H3K9La
Revert 700

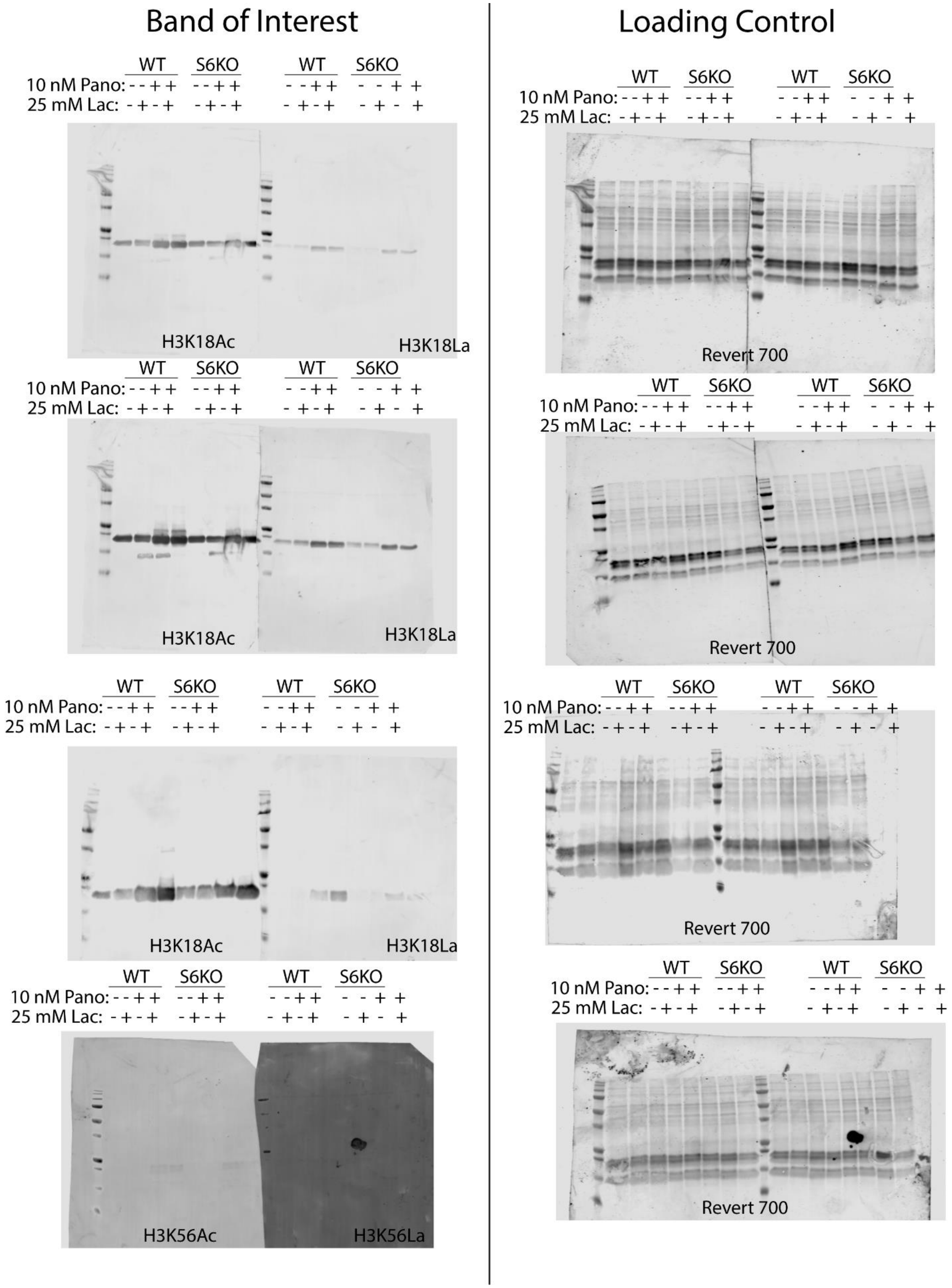
Band of Interest
Loading Control
WT S6KO WT S6KO
10 nM Pano: - - + + - - + + - - + + - - + +
25 mM Lac: - + - + - + - + - + - + - + - +
H3K18Ac
H3K18La
H3K56Ac
H3K56La
Revert 700

**Sirtuin 6 is a histone lysine delactylase (Supporting Information)**

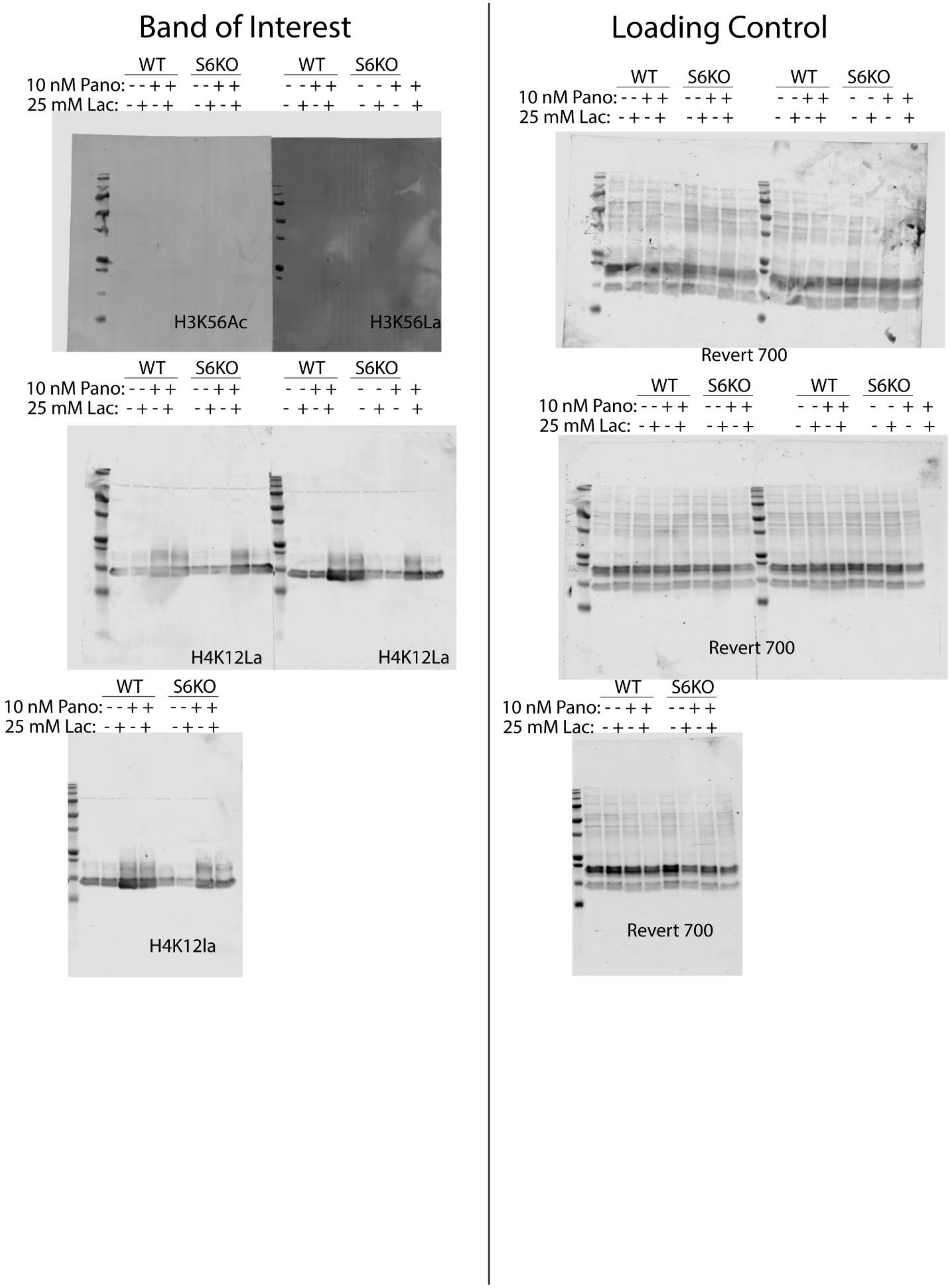

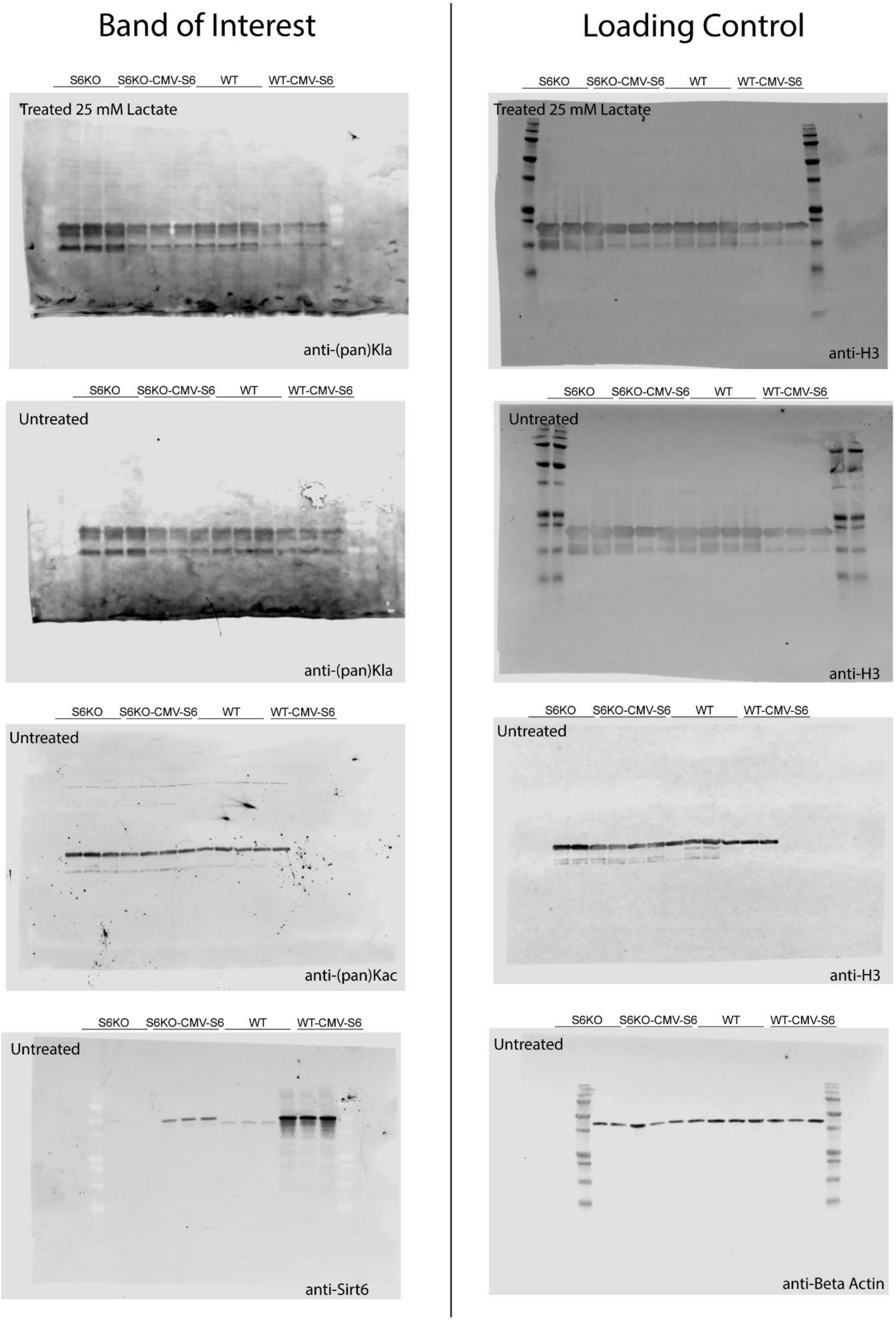
Band of Interest
Loading Control
S6KO S6KO-CMV-S6 WT WT-CMV-S6
Treated 25 mM Lactate
anti-(pan)Kla
S6KO S6KO-CMV-S6 WT WT-CMV-S6
Treated 25 mM Lactate
anti-H3
S6KO S6KO-CMV-S6 WT WT-CMV-S6
Untreated
anti-(pan)Kla
S6KO S6KO-CMV-S6 WT WT-CMV-S6
Untreated
anti-H3
S6KO S6KO-CMV-S6 WT WT-CMV-S6
Untreated
anti-(pan)Kac
S6KO S6KO-CMV-S6 WT WT-CMV-S6
Untreated
anti-H3
S6KO S6KO-CMV-S6 WT WT-CMV-S6
Untreated
anti-Sirt6
S6KO S6KO-CMV-S6 WT WT-CMV-S6
Untreated
anti-Beta Actin

**Sirtuin 6 is a histone lysine delactylase (Supporting Information)**

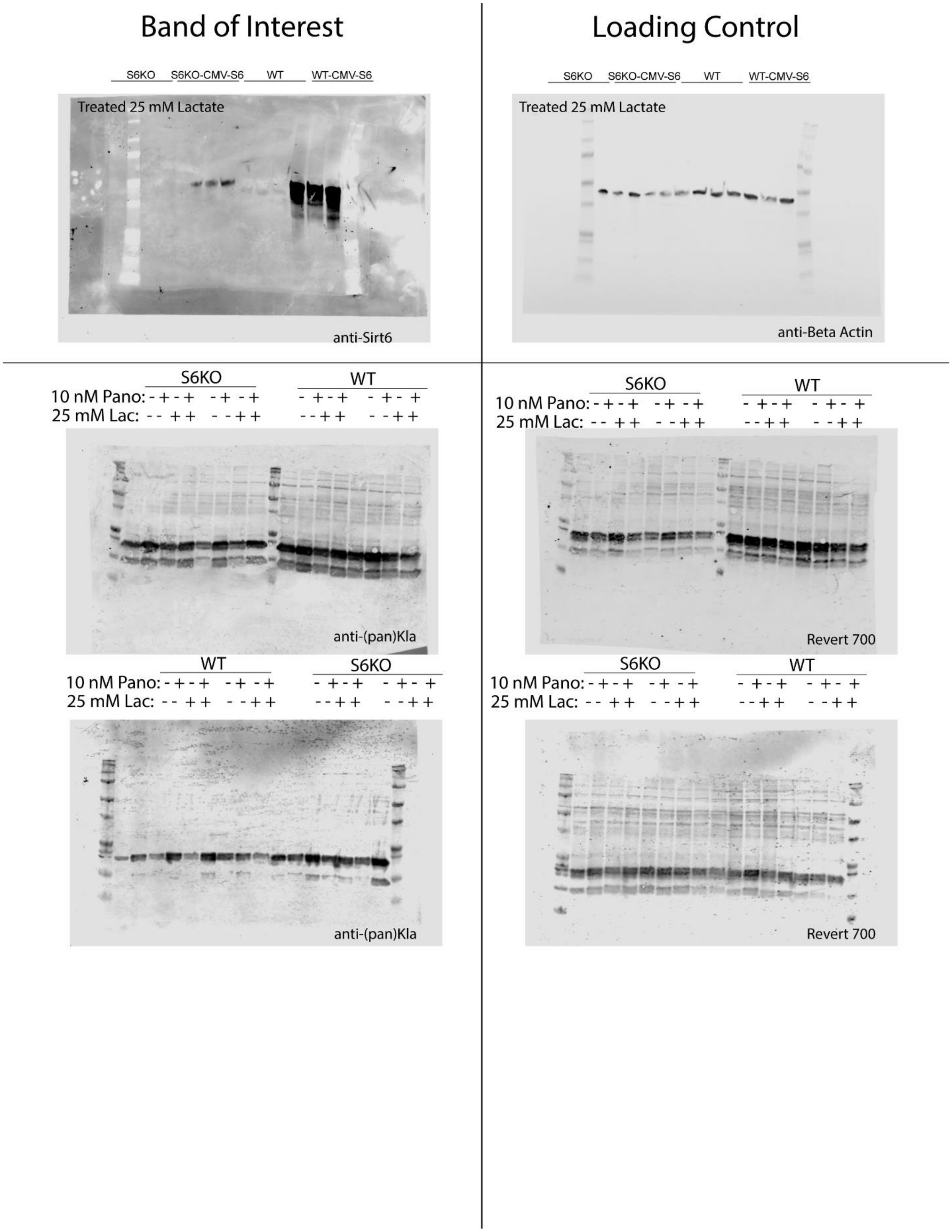

**Sirtuin 6 is a histone lysine delactylase (Supporting Information)**

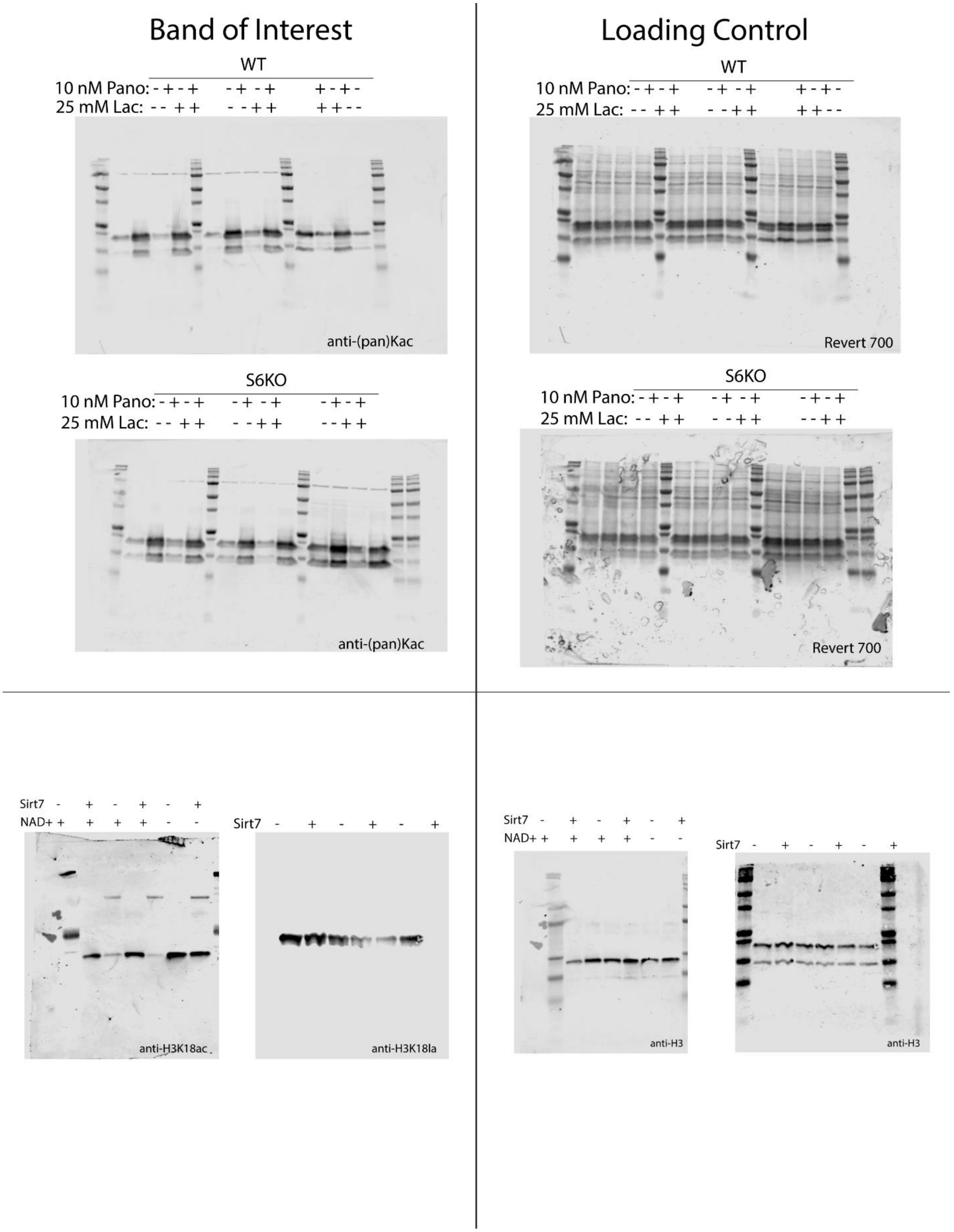

**Sirtuin 6 is a histone lysine delactylase (Supporting Information)**

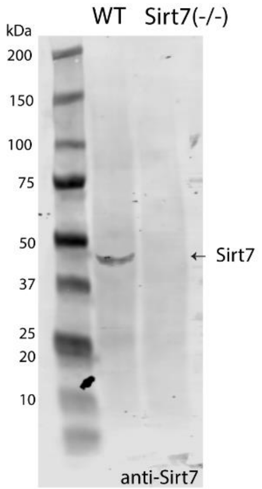


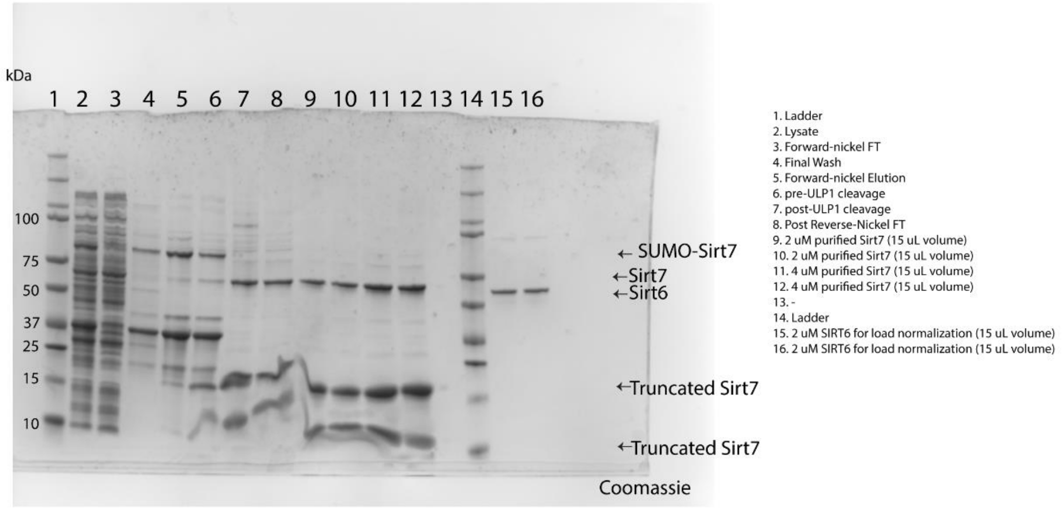


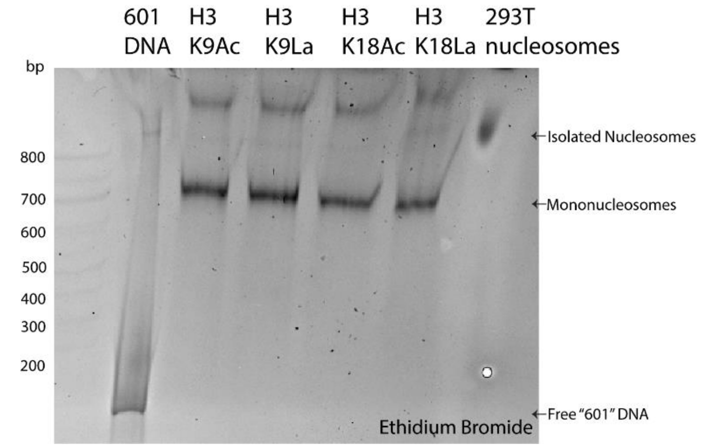
